# DASH/Dam1 complex mutants stabilize ploidy by weakening kinetochore-microtubule attachments

**DOI:** 10.1101/2022.09.28.509901

**Authors:** Max A. B. Haase, Guðjón Ólafsson, Rachel L. Flores, Emmanuel Boakye-Ansah, Alex Zelter, Miles Sasha Dickinson, Luciana Lazar-Stefanita, David M. Truong, Charles L. Asbury, Trisha N. Davis, Jef D. Boeke

**Author notes:** Equal Contribution (Guðjón Ólafsson, Rachel L. Flores).

## Abstract

Forcing budding yeast to chromatinize their DNA with human histones manifests an abrupt fitness cost. We previously proposed chromosomal aneuploidy and missense mutations as two potential modes of adaptation to histone humanization. Here we show that aneuploidy in histone-humanized yeasts is specific to a subset of chromosomes, defined by their centromeric evolutionary origins, however, they are not adaptive. Instead we show that a set of missense mutations in outer kinetochore proteins drive adaptation to human histones. Further, we characterize the molecular mechanism of two mutants of the outer kinetochore DASH/Dam1 complex, which reduce aneuploidy by suppression of chromosome instability. Molecular modeling and biochemical experiments show that these two mutants likely disrupt a conserved oligomerization interface thereby weakening microtubule attachments. Lastly, we show that one mutant, *DAD1*^E50D^, while suppressing chromosome instability in mitosis, leads to gross defects in meiosis. In sum, our data show how a set of point mutations evolved in the histone-humanized yeasts to counterbalance human histone induced chromosomal instability through weakening microtubule interactions, eventually promoting a return to euploidy.

## Introduction

Evolution is punctuated by moments of turmoil followed by rapid adaptation and stasis (Gould and Eldredge, 1993). Genomic studies have revealed a mode of “creation by crisis”, with mechanisms ranging from whole genome duplication in yeasts to chromothripsis and telomere crisis in cancer cells (Baca et al., 2013; Counter et al., 1992; Wolfe and Shields, 1997). These episodic bursts of innovation typically play out at the level of large-scale DNA mutation (Heasley et al., 2021). Cross-species genome hybridization via molecular engineering presents a dramatic new example of lab-directed genetic crisis (Boonekamp et al., 2022; Kachroo et al., 2015; Laurent et al., 2020). Because histone proteins are intimately associated with DNA, the complete exchange of native histone genes with non-native histone genes poses a substantial genetic barrier. We have previously shown that the budding yeast, *Saccharomyces cerevisiae*, can subsist with human histone comprised chromatin (Truong and Boeke, 2017). However, a large bottleneck limits how readily this occurs, with just ~1 in 10^9^ cells surviving the initial histone swap (Haase et al., 2019). The histone-humanized cells are initially extremely unfit, with a generation time ranging from 8-12 hours, but they quickly adapt and fitness improves. Adaptation is associated with the acquisition of distinct bypass mutations and accumulation of aneuploid chromosomes, but how these processes contribute to improved fitness is unclear. However, the high levels of chromosome instability suggest critical defects to machinery responsible for chromosome segregation in histone humanized strains.

Accurate segregation of chromosomes relies on ensuring proper centromere-kinetochore-microtubule connections. After replication, sister chromosomes must be captured by the spindle microtubules, bioriented, and segregated equally into daughter cells (Nicklas, 1997). The establishment of kinetochore biorientation is the critical step in ensuring faithful chromosome segregation. Failure to establish correct orientation causes chromosome missegregation and aneuploidy, which leads to decreased cellular fitness in lab yeasts (Hose et al., 2020; Torres et al., 2007, 2010) and underlies many human maladies (Antonarakis, 2017; Oromendia and Amon, 2014).

Centromeres serve as the coupling point of chromosomes to the spindle microtubules – an interaction which is bridged by the megadalton kinetochore complex (Biggins, 2013). The centromeric variant histone H3 (Cse4 in *S. cerevisiae*, CENP-A in humans) plays a central role in the process of chromosome segregation by defining the region of centromeric DNA in the majority of species (Steiner and Henikoff, 2015). In *S. cerevisiae*, centromeres are defined as a single Cse4-containing nucleosome that wraps a specific sequence of ~125 base pairs (bp) of DNA. Coupling to a single microtubule is achieved through the association of a Cse4-containing nucleosome with the inner kinetochore protein complexes CCAN and Cbf3 (Biggins, 2013; Cottarel et al., 1989; Furuyama and Biggins, 2007; Winey et al., 1995). From here adaptor complexes, MIND^MIS12^ and CNN1^CENP-T^, bridge the gap to link to the outer kinetochore complexes Ndc80c and DASH/Dam1c that interface with microtubules (Jenni et al., 2017).

Kinetochore-microtubule attachments made by Ndc80c and DASH/Dam1c are highly regulated to ensure incorrect attachments are not over stabilized (Tien et al., 2010). Directed destabilization of incorrect attachments allows for attachments to be released and corrected. This regulation is achieved through kinetochore-microtubule turnover driven by the kinase activity of Aurora B kinase (Carmena et al., 2012; Cimini et al., 2006; Pinsky et al., 2006; Tanaka et al., 2002). Aurora B (Ipl1) forms the chromosomal passenger complex (CPC), alongside of INCENP (Sli15), Borealin (Bir1), and Survivin (Nbl1), whose recruitment to centromeric chromatin stimulates correction of incorrect microtubule attachments (Carmena et al., 2012; Kawashima et al., 2010; Yamagishi et al., 2010). As these mechanisms of CPC recruitment involve the direct interaction with nucleosomes (Abad et al., 2019), it is thus plausible human histones may disrupt this pathway and lead to chromosome instability in histone humanized yeasts.

Whole genome sequencing of histone-humanized yeasts hinted towards mutation of outer kinetochore genes and aneuploidy as two potential paths of adaptation (Truong and Boeke, 2017). Here we set out to answer the relative contributions of mutation and aneuploidy to adaptation to human histones. We find that aneuploidy is all together non-adaptive and that aneuploidy accumulation is biased to non-random subset of chromosomes based on their centromeric evolutionary origins. Instead, using genetic, molecular modeling, and biochemistry techniques we show that a set of DASH/Dam1c mutants are adaptive to yeasts with human histones. Together our data support a mechanism whereby DASH/Dam1c mutants disrupt their own oligomerization, which weakens kinetochore-microtubule attachments thereby suppressing chromosome instability and reducing the incidence of aneuploidy.

## Results

### DASH/Dam1c mutants are dominant suppressors of human histones

Histone-humanized yeasts are generated by using a low background dual-plasmid histone shuffle assay followed by counterselection of the yeast histone genes, which are linked to the *URA3* marker, with 5-FOA (Figure 1A, S1A; (Haase et al., 2019; Truong and Boeke, 2017). After approximately 2 - 4 weeks of growth, we observed candidate humanized colonies as indicated by the appearance of small colonies, whereas larger colonies retained yeast histone genes (Figure S1B). We validated eight new lineages (yHs9-16) as bona fide humanized clones by the loss of the yeast histone genes as determined by PCR analysis (Figure S1B) and whole genome sequencing (Figure S1C).

**Figure 1.**
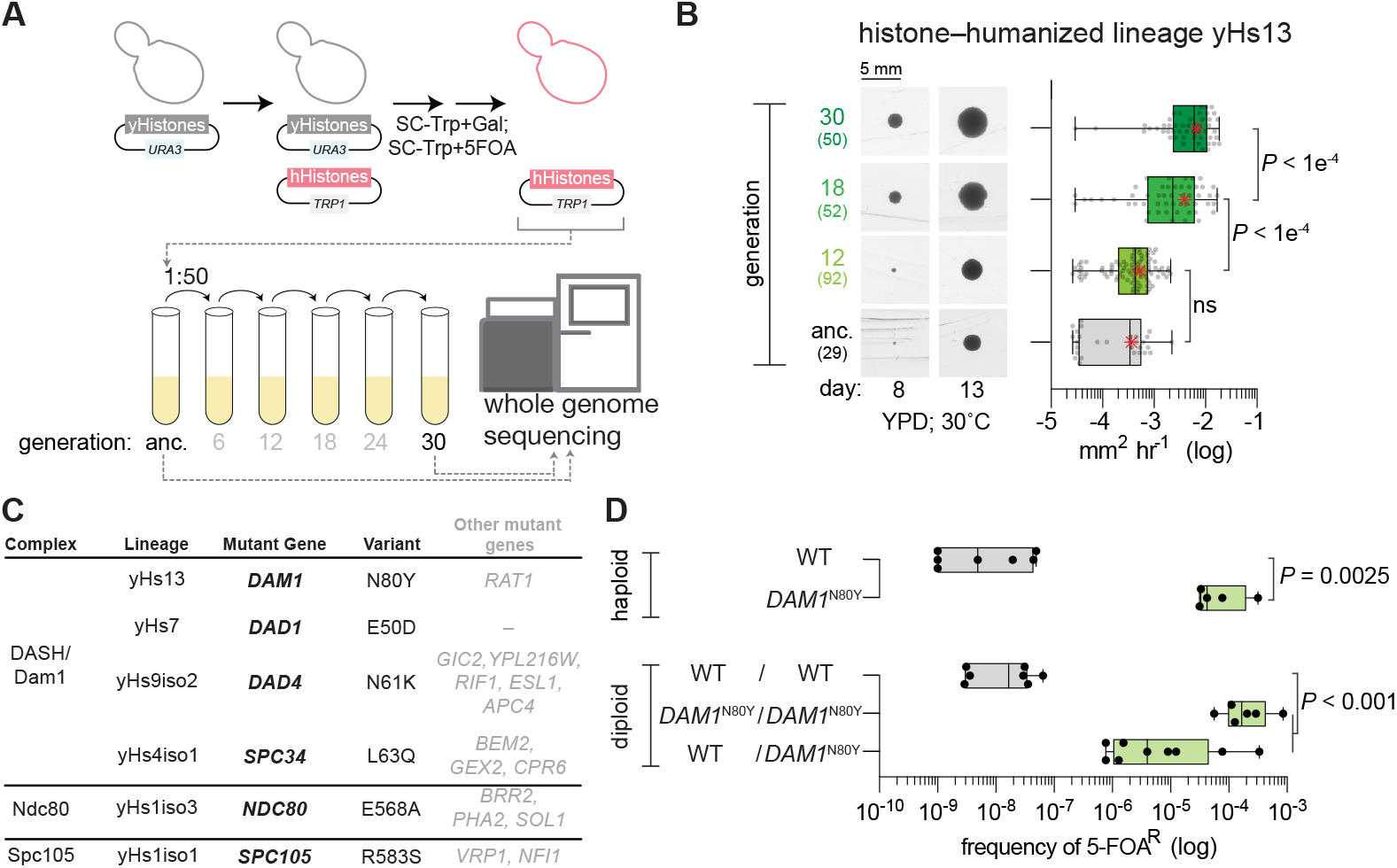
DASH/Dam1c mutants are dominant genetic suppressors of histone humanization. (**a**) Dual-plasmid histone shuffle strategy used to generate histone-humanized yeasts (see methods for details). Humanized isolates were passaged in rich medium for at least 5 cycles, at which point the ancestral isolate and evolved were sequenced. (**b**) Growth assay of ancestral and evolved histone-humanized lineage yHs13. Cells from the ancestral populations and evolved populations (indicated by generations in rich medium) were restruck onto rich medium agar plate and colonies were imaged for up to three weeks to observed the change in colony size (left images). The average growth rate in mm^2^ hr^-1^ was calculated by taking the change in colony size between time points divided by the time interval (right graph). Numbers in parentheses indicates the number of colonies analyzed. Significance of mean differences in growth rates was determined with an ordinary one-way ANOVA multiple comparisons with Turkey correction of multiple hypothesis tests. (**c**) Table of histone–humanized lineages that evolved a mutation in an outer kinetochore complex. Variant column shows the observed nonsynonymous alteration of the bolded gene in the mutant gene column. Other genes which were mutated in each lineage are shown in non-bolded font, details of these mutations can be found in supplemental table 3. (**d**) Sufficiency validation for the *DAM1*^N80Y^ mutation demonstrates the *DAM1*^N80Y^ mutation significantly increases the rate of humanization over wild type (Two-tailed unpaired t test; *p* = 0.0093). Green filled bars indicate successful isolation and confirmation of humanized yeasts. Significance of the mean difference in 5-FOA^R^ frequency was determined with the Mann-Whitney test.

To see how these lineages adapt to human histones we continually passaged them in rich medium and observed a dramatic fitness improvement within a brief period of 30 generations (corresponding to five passages lasting a total of approximately 25 days; Figure 1A, S1D). For example, in the histone-humanized lineage yHs13 the mean growth rate significantly improved over 17 fold between the ancestral and the generation 30 descendant (Figure 1B). We found that lineage yHs13 initially evolved a mutation in the DAM1 gene – a component of the outer kinetochore complex DASH/Dam1c (Figure 1C). Additionally, we identified 60 unique nucleotide variants from the ancestral and evolved lineages (Supplemental tables 1 and 2), more than doubling the total number of candidate suppressors of histone humanization from previously reported (Truong and Boeke, 2017). On average each strain had 3 mutations (Figure S1E) and we observed only two genes, *URA2* and *GEX2*, with more than one mutation (Truong and Boeke, 2017) and Supplemental table 1). This set of genes was biased toward processes related to the cell cycle, chromosome segregation, rRNA processes, and chromatin remodeling (Figure S2). Across all of our histone–humanized lineages we isolated six independent lineages with mutations in the outer kinetochore protein complexes DASH/Dam1c, Ndc80c, and Spc105c, suggesting that alteration to microtubule attachments may be a potent route for adaptation (Figure 1C).

The DASH/Dam1c mutations arose in lineages with additional mutations (Figure 1C). We, therefore tested the sufficiency of these mutations in suppressing the fitness defect associated with histone humanization. First, we used CRISPR-Cas9 mediated mutagenesis to scarlessly introduce each of the non-synonymous mutations into an isogenic haploid histone shuffle strain, and then derived both heterozygous and homozygous diploid histone shuffle strains by mating (Figure S3A-B, Table S1). The majority of mutants had no growth defects in the wild-type (WT) background, except for a general growth defect of the *dad4*^N61K^ mutant and cold sensitivity at 4°C for the *DAM1*^N80Y^ mutant (Figure S3C, where dominant mutations appear capitalized).

Sufficiency of each mutation was tested using our histone plasmid shuffle assay (Figure 1A, Figure S1A). We defined sufficiency as the ability of a mutant to robustly generate histone-humanized colonies that, upon counter-selection of the yeast histone plasmid with 5-FOA, are resistant to 5-FOA (5-FOA^R^) and devoid of yeast histones. For example, in a haploid strain, the *DAM1*^N80Y^ mutant increased the frequency of 5-FOA^R^ over ~4,000 times that of the WT (Figure 1D). Likewise, the homozygous diploid *DAM1*^N80Y^ mutant generated 5-FOA^R^ colonies at ~10,000 times the diploid WT frequency. Further, we found that *DAM1*^N80Y^ is dominant to WT, as the heterozygous diploid strain produced 5-FOA^R^ colonies at a frequency ~3,000 times higher than the diploid WT (Figure 1D). We obtained similar results for all tested mutants except for dad4^N61K^, which only weakly humanized in the heterozygous diploid background (Figure S3D-E; all calculated humanization frequencies can be found in Supplemental table 4). Importantly, these mutants lead to faster growth with humanized colonies that appear within 14 days upon counterselection (Figure S3D), in contrast to the WT background, where colonies can take over 21 days to first appear (Truong and Boeke, 2017). These data show that the DASH mutants are dominant and significantly increased frequency of generating histone humanized yeasts. We next sought to disentangle the contributions of aneuploidy and the DASH mutants to adaptation to human histones.

### Ancient paralogous centromere origins best explain aneuploid frequency

One consistent feature in all ancestral humanized lineages is the presence of aneuploid chromosomes (Figure 2A, Figure S4, Supplemental table 3; Truong and Boeke, 2017). In fact, the majority of the lineages maintained their aneuploidies even as their fitness improved (Figure S1D, Figure S4). The aneuploids occurred nonrandomly, with chromosomes *I, II, III*, and *XVI* being the most frequently aneuploid (Figure 2A). We examined a number of candidate chromosomal features but found the frequency of aneuploidy in histone humanized yeast did not strongly correlate with any of them (Figure S5A). Two features showed borderline significance, as we observed a negative correlation with chromosome size (Pearson r = −0.54; *p* = 0.037) and a positive correlation with centromere AT percentage (Pearson r = 0.51; p = 0.044), however these features were not necessarily predictive of aneuploidy (Figure S5A-C). We were thus curious if any additional chromosome feature may explain the observed frequency of aneuploidy in histone humanized yeasts.

**Figure 2.**
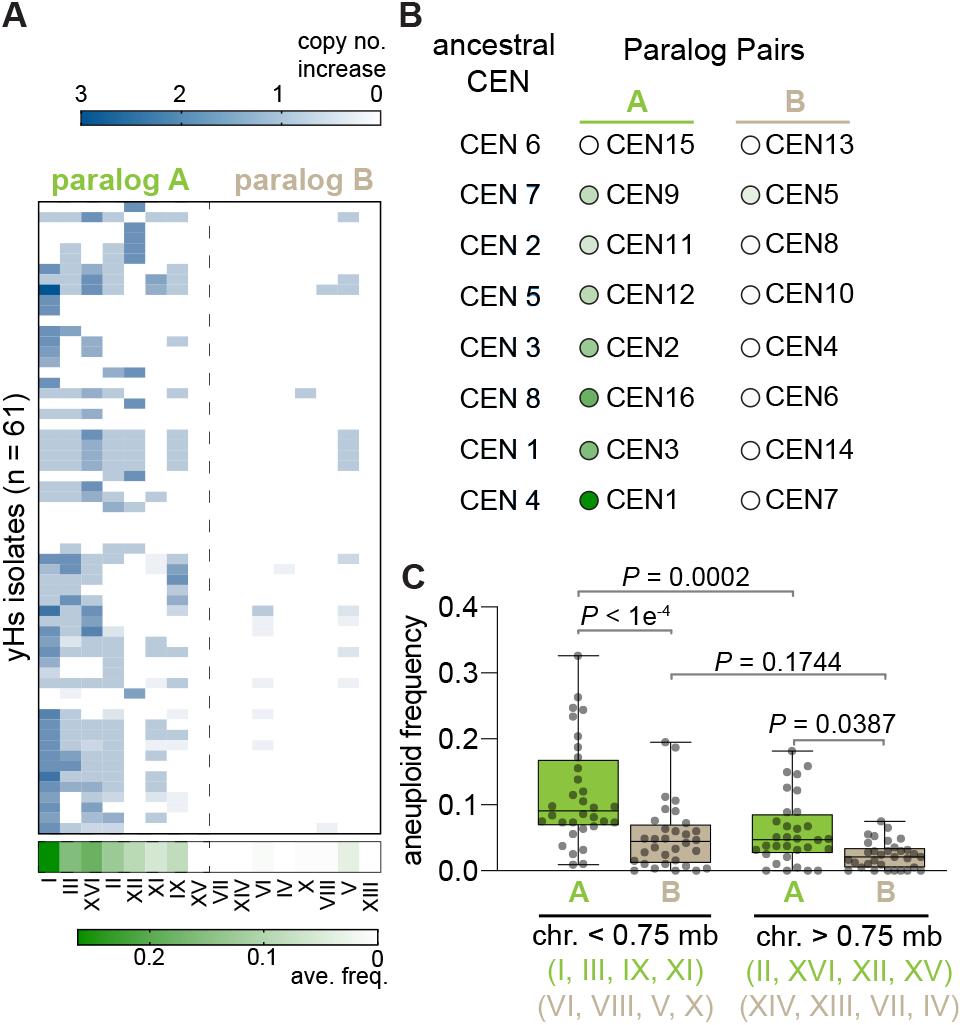
Ancient paralogous centromere pairs best explain aneuploid frequency. (**a**) Aneuploid frequency in histone–humanized yeasts. Chromosomes are displayed divided into two groupings based on paralogs (i.e., Chr I is the paralog of Chr VII). The average frequency of aneuploidy for each chromosome is shown on the bottom in the green colored heat map. Each row represents the aneuploidy for a single humanized lineage either from this study and Truong and Boeke, 2017. (**b**) Difference in aneuploid frequency between paralog pairs, colored circles next to chromosomes indicate the average aneuploid frequency in histone humanized yeast as defined in panel A. Paired t-test between the mean aneuploid frequency of paralog group A and B p = 0.004. Ancestral *CENs* labeling are taken from Gordon et al., 2011. (**c**) Comparison of the frequency of aneuploidy in yeasts based on two factors: chromosome size (8 smallest chromosomes; blue (*VI, I, III, IX, VIII, V, XI, X*) – less than 0.75 Mb vs. 8 largest; red (*XIV, II, XIII, XVI, XII, XV, VII, IV*) – greater than 0.75 Mb) and centromere paralog group (coloring indicates paralog group as in panel A). Aneuploid frequencies are from our study plus seven additional studies that reported aneuploidy across a variety of yeast strains (see main text). Significance difference of the mean frequency of aneuploidy was tested with ordinary one-way ANOVA multiple comparisons with Turkey correction of multiple hypothesis tests.

The extant karyotype of *S. cerevisiae* is the product of an allopolyploidization event occurring ~93 million years ago (Marcet-Houben and Gabaldón, 2015; Shen et al., 2018; Wolfe and Shields, 1997) – where the lineage leading to *S. cerevisiae* maintained all eight paralogous pairs of centromere sequences (Gordon et al., 2011). It is striking that eight chromosomes displayed frequent aneuploidy and eight did not (Figure 2A), thus, we hypothesized that the observed differences in aneuploid frequency might be a reflection of differences between centromeric paralogs. The list of centromere paralog pairs, as determined by synteny analysis (Gordon et al., 2011), is displayed in figure 2B. We divided paralogs from each pair into group A or group B, where group A centromeres have higher frequency of aneuploidy in histone humanized yeast than those in group B. We note that one paralog pair *CEN15* - *CEN13* both presented zero aneuploidies in our data set, thus we excluded the pair from subsequent analysis (Figure 2A-B). To see if the bias in aneuploid frequency between group A and B was a trend observed in other studies, we analyzed the frequency of chromosomal aneuploidy across another seven studies (Duan et al., 2018; Gallone et al., 2016; Kao et al., 2010; McCulley and Petes, 2010; Peter et al., 2018; Sharp et al., 2018; Zhu et al., 2016). This data set represents a total of 1,391 occurrences of aneuploids across 1,688 yeast strains from a variety of sources (i.e., wild strains, industrial strains, clinical strains, and laboratory strains; Supplemental table 5). We found that the paralog pairs displayed a bias in the frequency of aneuploid, with group A showing higher frequency of aneuploidy across the unrelated studies (Figure S5D; paired t-test between mean difference in paralog pair’s aneuploid frequencies p = 0.0001846). We next asked how frequently random pairings of the same data would lead to such extremely different aneuploidy frequencies between chromosome pairs (in other words, how extreme is the difference between group A and group B pairs when compared to other combinations of groupings). To this end, we performed randomization tests to make random reallocations of the aneuploid counts from all eight studies into two groupings. This allowed us to create a distribution of *t*-statistics of the mean difference assuming the null hypothesis was true (i.e. there is no difference in aneuploid frequency between the two groups; Figure S5E). We find that the observed difference in aneuploid frequency between group A and B is significantly greater than chance alone would predict, supporting our hypothesis that centromere paralog type (A or B) is leading to a non-random aneuploidy landscape (Figure S5E; p < 0.00004). Lastly, we asked if chromosome size effected the rate of aneuploidy. We observed that the frequency of aneuploidy for chromosomes in the paralog group B is significantly lower than for the chromosomes in paralog group A, even when controlling for chromosome size (Figure 2C). Further, there is not a significant difference in aneuploid frequency within group B chromosomes when grouping chromosomes by size (Figure 2C). Additionally, the number of aneuploids per chromosomes showed a significant negative correlation with chromosome size when considering only the chromosomes as distinct groupings (Figure S5F), suggesting that group B chromosomes have unexpectedly lower frequency of aneuploidy than chromosome size alone would predict. In sum, these data suggest that yeast have a non-random aneuploidy landscape, which is responsible for the observed frequency of aneuploidy in our data.

### Chromosomal aneuploidies are not adaptive to life with human histones

The nonrandom aneuploidy landscape driven by paralog type, predicts aneuploidy is non-adaptive in histone humanized yeasts, we therefore explicitly test this prediction. For this experiment we used the histone humanized strain yHs5, which has the scc4^D65Y^ mutation and has eight persistently aneuploid chromosomes (Chromosomes *I, II, III, V, IX, XI, XII, XVI;* Truong and Boeke (2017)). We hypothesized that if aneuploidy were adaptive, then a strain with preexisting aneuploidy would generate histone humanized yeast at a higher rate than isogenic a euploid strain. To generate the isogenic strains we “captured” yHs5 clones in various ploidy states by “re-yeastification” of its histones. This was achieved by transforming yHs5 with a plasmid encoding yeast histones (Figure S6A). We assessed twelve “re-yeastified” transformants for aneuploidy using flow cytometry and observed that eight transformants had heavy aneuploid loads and four transformants were either euploid or near-euploid (Figure S6B). Our prediction, if aneuploidy were adaptive, was that when we humanized these strains again the ‘re-yeastified’ strains with preexisting aneuploids would humanize at a higher frequency than the euploid counterparts. On the contrary, we observed that the level of preexisting aneuploidy did not affect the humanization rate at all, as both isogenic aneuploid and euploid shuffle strains humanized the same rate (Figure S6C). From this experiment we conclude that aneuploidy does not provide a selective advantage for histone-humanization. These data are consistent with the idea that the missense scc4^D65Y^ mutation is the driving force for adaptation to histone humanization. This suggests the observed frequency of aneuploidy in histone humanized yeasts is a consequence of a non-random aneuploidy landscape, in which paralog group A chromosomes are sensitized to human histone induced chromosome instability.

### Histone-humanized yeasts display chromosome instability caused by centromere dysfunction

The above results suggests that human histones induce chromosome instability. Our previous work showed that centromeric DNA is sensitized to micrococcal nuclease (MNase) digestion in the histone-humanized yeasts (Truong and Boeke, 2017). This is consistent with lower occupancy of the centromeric nucleosome. Normally, the 16 centromere-kinetochore attachments are clustered throughout the entirety of the cell cycle and transcriptional activity at centromeres, which is tightly regulated, is important for establishing these attachments (Hedouin et al., 2022; Ling and Yuen, 2019). We reasoned that human histones may expose centromeric DNA and create more transcriptional open chromatin. We first, reanalyzed MNase digested chromatin sequencing data to infer nucleosome occupancy. We observed that centromeric nucleosome occupancy is severely depleted in histone-humanized yeasts (Figure 3A). This was true of both humanized lineages tested (yHs7 and yHs5, with the *DAD1*^E50D^ and scc4^D65Y^ mutations, respectively), with no significant differences between the two genotypes.

**Figure 3.**
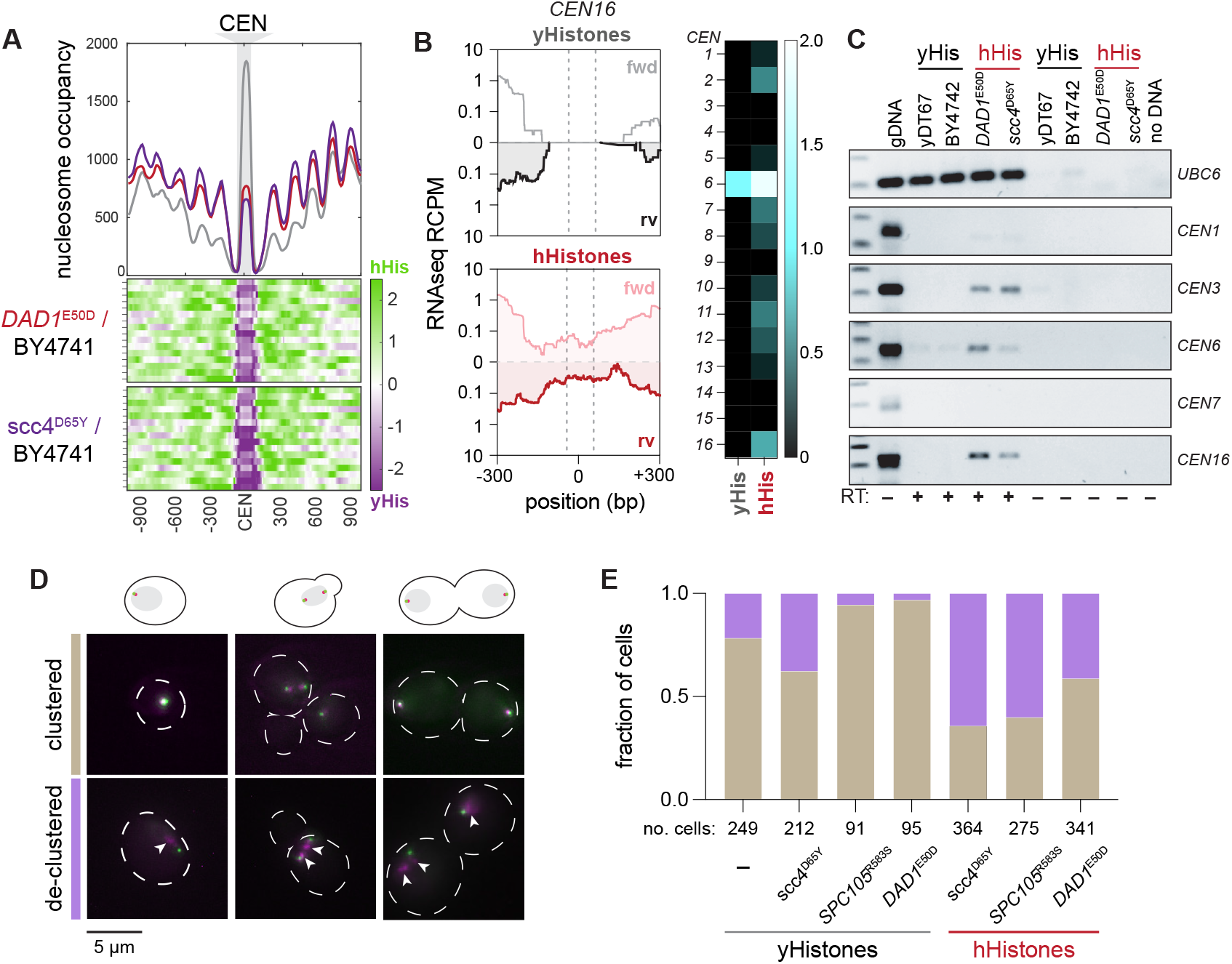
Histone humanization cause centromere dysfunction in yeast. (**a**) Centromeric nucleosome occupancy. The relative nucleosome occupancy plots are given for each strain in the top plot (grey, wildtype; red, *DAD1*^E50D^ humanized; purple, scc4^D65Y^ humanized). Values represent the chromosome mean occupancy 900 bp up and downstream of all 16 centromeres (each centromere occupancy values being an average of 3 biological replicates). Bottom plots, log2 ratio nucleosome occupancy enriched in humanized (green) and depleted in humanized (purple), each row represents one of the sixteen centromeric regions. (**b**) Total RNA sequencing of cenRNAs in yeasts. Left; read counts per million tracks for *CEN16* in wildtype (black, top) and *DAD1*^E50D^ humanized (red, bottom) yeasts. Right; heatmap of cenRNAs transcripts per million counts across all chromosomes in wildtype and *DAD1*^E50D^ humanized strains. Centromere *VI* is marked with an asterisk to indicate that the plasmid encoding the histone plasmid also encodes a *CEN6* sequence, which reads likely emanant from. (**c**) RT-PCRs of the indicated RNAs. Lane 1 is genomic DNA, lane 2 is wildtype shuffle strain (yDT67), lane 3 is BY4742, lane 4 is *DAD1*^E50D^ humanized, lane 5 is scc4^D65Y^ humanized, lanes 6–9 are the same as 2–5, but without reverse transcriptase added, lane 10 is no DNA. (**d**) Example images from strains with fluorescent tags on the microtubule organizing center (Spc110) and kinetochore (Nuf2). (**e**) Bar plots of the fraction of cells with clustered Nuf2-RFP foci. Brown shading indicates fraction of cells with correct clustering, and the purple shading indicates fraction of cells with declustered Nuf2-RFP foci.

We next assessed the levels of CEN transcription genomewide by total-RNA sequencing on WT (those with yeast histones) and histone humanized yeasts. Since CEN RNAs are rare in WT yeast, this assay is expected to fail to detect any meaningful amount of CEN RNAs (Hedouin et al., 2022). In our data we were able to identify robust CEN RNA transcription from the humanized lineages tested, while failing to detect such transcription in WT cells (Figure 3B–C). *CEN6* RNA was the most robustly transcribed centromere (Figure 3B; right). This result can be attributed to reads that emanate from the plasmid-borne centromere sequence - plasmids encoding the histone genes also encode a minimal *CEN6* sequence - and not to the native chromosomal location. Indeed we observed fewer reads mapping to chromosomal *CEN6* in WT cells versus humanized cells (Figure S7). These results were confirmed by RT-PCR on a subset of yeast CEN RNAs (Figure 3C). Importantly we used a polyT^18^ oligo for the RT reaction as CEN RNAs are polyadenylated (Ling and Yuen, 2019). The two humanized lineages tested showed robust CEN RNAs expression for chromosomes *III*, *VI*, and *XVI*, whereas the two WT controls showed little to no expression (Figure 3C). Given that, the RT-PCR primers used do not complement to the plasmid-borne *CEN6* sequence, we observed little amplification in the WT strains. In sum, these results suggest that elevated CEN RNA transcription is a general defect caused by human histones. Altogether our results indicate that the structure of the histone-humanized centromeric chromatin is under a persistent state of dysfunction, as indicated by the increased MNase sensitivity and transcription of the centromeres.

Given the strong functional relationship between centromere and kinetochore, we wondered if their coupling could also be compromised in the histone humanized yeasts. We next investigated kinetochore clustering by imaging log phase cells with a RFP-tagged kinetochore protein Nuf2 in WT and humanized backgrounds (Figure 3D). For these first experiments we used three human histone suppressor mutations, *DAD1*^E50D^, *SPC105*^R583S^, and scc4^D65Y^, the latter two represent control genotypes for euploid vs. aneuploid (scc4^D65Y^) and for outer kinetochore vs. specifically DASH/Dam1c (SPC105^R583S^). In strains with yeast histones, we observed moderate levels of Nuf2-RFP foci declustering in both WT (note, WT here is the histone shuffle strain) and scc4^D65Y^ stains, however, for both the *DAD1*^E50D^ and SPC105^R583S^ mutants with yeast histones we observed improved Nuf2-RFP clustering (Figure 3E).

We next generated histone-humanized strains with the Nuf2-RFP for the three mutants and performed the same imaging experiment. In our three histone-humanized strains, we observed substantial Nuf2-RFP declustering, with over 70% of cells in the *scc4*^D65Y^ background exhibiting declustered Nuf2-RFP foci (Figure 3E). This data is consistent with centromere dysfunction driving higher rates of chromosome instability. Intriguingly, the *DAD1*^E50D^ background showed only 30% of cells with declustered Nuf2-RFP foci, whereas SPC105^R583S^ background showed over 60% with declustered Nuf2-RFP foci. Remarkably, the humanized *DAD1*^E50D^ mutant showed little difference with the scc4^D65Y^ mutant in terms of CEN RNA transcription or CEN MNase sensitivity, in spite of its marked improvement of kinetochore clustering (Figure 3A–C,E). This suggest that *DAD1*^E50D^ mutant, and perhaps other DASH mutants, rescues chromosome instability independently of directly rescuing the human histone induced centromere dysfunction.

### DASH/Dam1c mutants suppress chromosome instability and lead to euploidy

As we have shown above aneuploidy in histone humanized yeasts is likely caused by non-random chromosome instability initiated by human histone induced centromere dysfunction. We were curious if the DASH mutants suppress chromosome instability, as suggested by the kinetochore clustering data. We therefore sought understand the dynamics of aneuploidy from the ancestral to the evolved histone humanized strains in lineages that evolved DASH mutations. Strikingly, the lineage yHs13, which evolved the *DAM1*^N80Y^ mutation, progressed from presenting chromosome XII aneuploidy to complete euploidy in the evolved strain (Figure 4A). Across all lineages sequenced, we identified a total of four lineages (yHs4, yHs7, yHs9, and yHs13) with mutant DASH/Dam1c genes (Supplemental table 2 and Truong and Boeke, 2017), and on average these evolved lineages had significantly fewer aneuploid chromosomes compared to all other evolved lineages (Figure 4B). Furthermore, these four lineages saw a decrease, on average, of 2 to 3 fewer aneuploid chromosomes when comparing the evolved to the ancestral isolates (Figure 4C). These results suggest that the DASH mutants suppress chromosome instability and have an “aneuploidy-reducing” phenotype.

**Figure 4.**
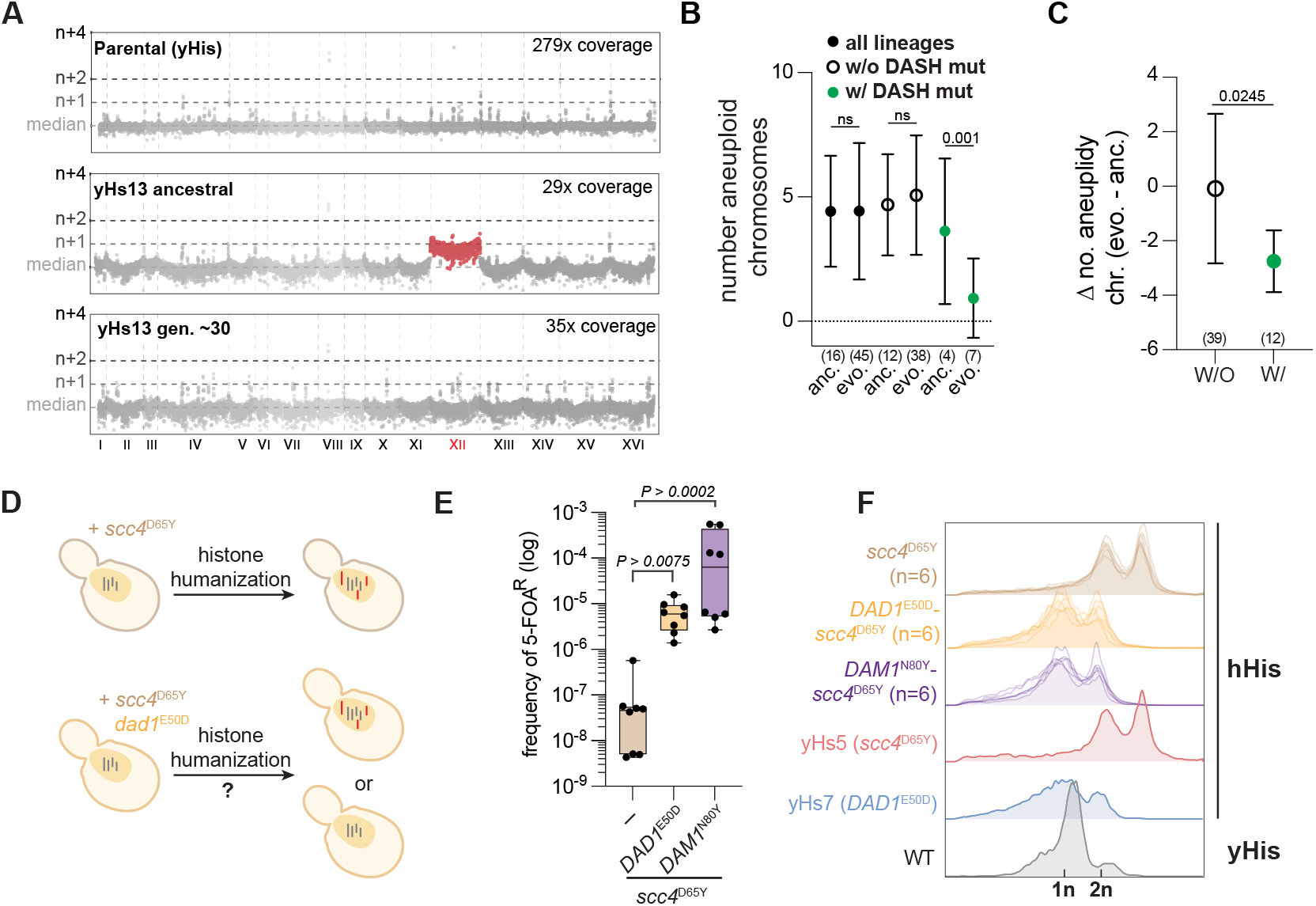
DASH/Dam1c mutations are sufficient for aneuploid reduction. (**a**) Genome-wide chromosome coverage maps for the parental (yeast histones) and the ancestral and evolved yHs13 strains (note the loss of aneuploid chromosome XII in the evolved isolate). Note that yHs13 is the lineage that evolved the mutation *DAM1*^N80Y^. Chromosome coverage copy was normalized as the log2 ratio of the average coverage of a 1 Kb window by the genome median coverage. (**b**) Evolution of aneuploid chromosomes in histone-humanized strains. The number of aneuploid chromosomes is given for each of the indicated grouping of humanized strains. Sample size is given below each (number of histone–humanized lineages), Dunnett’s tests of the mean differences between the three ancestral-evolved pairs are provided. (**c**) Change in the number of aneuploid chromosomes in humanized lineages with or without DASH/Dam1c mutations. Sample size is given below each, unpaired t-test of the mean difference is provided. (**d**) Aneuploidy accumulation assay. Histone–humanization in the scc4^D65Y^ genetic background results in persistent aneuploidy (red chromosomes), does the presence of a second mutation, *DAD1*^E50D^ or *DAM1*^N80Y^, stop the accumulation of aneuploidies to maintain euploidy (grey chromosomes)? (**e**) DASH mutants increase histone–humanization in the scc4^D65Y^ background. Humanization rates of the scc4^D65Y^ mutant with and without *DAD1*^E50D^ or *DAM1*^N80Y^. Significance of the mean difference in 5-FOA^R^ frequency was determined with the Mann-Whitney test. (**f**) Ploidy analysis by flow cytometry in the scc4^D65Y^ humanized mutants from panel d. Flow cytometry data from humanized lineages yHs5 and yHs7 are shown for aneuploid and euploid controls, respectively.

First, we performed a laboratory evolution experiment to test the sufficiency of the DASH mutants apparent “aneuploidy-reducing” phenotype (Figure S8A). To this end, we took an already histone–humanized lineage (yHs16), which harbored seven aneuploid chromosomes in the ancestral strain (Figure S8A, S9A), and transformed in a plasmid containing either DAD1^WT^ or the mutant *DAD1*^E50D^ sequence. Three transformants each were selected and continually passaged in media to select for the plasmid over the course of 4 months, totaling ~90 generations. The ancestral strains, and evolved populations at ~30 and ~90 generations were frozen and whole genome sequencing was performed to determine chromosomal copy number. The clones transformed with the DAD1^WT^ plasmid did not show a significant change in the copy number of the ancestral aneuploid chromosomes, highlighting the persistent nature of aneuploidy in humanized strains (Figure S8A-C, S9B). In contrast, 2 of the 3 clones transformed with mutant *DAD1*^E50D^ showed nearly complete loss of all aneuploid chromosomes by 90 generations. Notably, one clone achieved euploidy almost immediately after acquiring *DAD1*^E50D^, while the other more slowly reduced aneuploidies as the experiment progressed (Figure S8A-C, S9C). Interestingly, clone one, which immediately returned to euploidy, eventually lost the *DAD1*^E50D^ mutant plasmid at generation 90 (Figure S8D). Further, the loss of *DAD1*^E50D^ plasmid coincided with the appearance of two new mutations including Sli15^D331Y^, a component of the chromosomal passenger complex (CPC) that regulates kinetochore-microtubule attachments, and Smc5^H984N^, sub-unit of the Smc5-Smc6 complex involved in chromosome separation (Figure S8E-F). The return-to-euploidy in both clones was confirmed by computing the log2 ratio median chromosome coverage, which returned to a log2 genome-median of ~0 (Figure S8C). This experiment was conducted in the presence of the native genomic DAD1 gene, where the mutant gene - encoded on an episomal plasmid - is in the context of additional mutations (Supplemental table 2, see yHs16). Therefore, we cannot exclude the hypothesis of reduced phenotypic penetrance of *DAD1*^E50D^ in this experiment. These results suggest that at least some mutants of the DASH complex are sufficient for aneuploidy reduction.

We next asked if these DASH mutants could prevent aneuploid chromosomes from accumulating (Figure 4D). To assess this, we compared the humanization frequency of the scc4^D65Y^ mutant shuffle strain in the presence or absence of DASH mutants (DAD1^E50D^ and *DAM1*^N80Y^). While the rate of humanization for scc4^D65Y^ mutant alone was low (~1 - 10 per million cells), in the presence of either the *DAD1*^E50D^ or *DAM1*^N80Y^ mutants the humanization rate increased 100 - 1000-fold (Figure 4E). In the shuffle strain with only the mutation *scc4*^D65Y^, we observed concomitant increases of the ploidy levels upon humanization (Figure 4F). In stark contrast, when the combination of mutations of either *scc4*^D65Y^ with *DAD1*^E50D^ or *DAM1*^N80Y^ were humanized, neither clone showed evidence of aneuploidy, as measured by DNA content by flow cytometry (Figure 4F). Therefore, we conclude that both *DAD1*^E50D^ and *DAM1*^N80Y^ mutants are sufficient to suppress chromosome instability and cure aneuploidy in the histone–humanized yeasts.

### DASH/Dam1c mutants suppress ipl1-2 driven chromosome instability

The Aurora B kinase, IPL1 in yeast, is the master regulator of kinetochore-microtubule attachments, the disruption of which leads to defects in chromosome segregation (Biggins et al., 1999; Pinsky et al., 2006). We therefore examined the interaction with the temperature sensitive ipl1-2 allele and our DASH/Dam1 mutants. The mutant ipl1-2 is known to increase ploidy by chromosome missegregation, causes severe growth defects at non-permissive temperatures and sensitivity to benomyl (Chan and Botstein, 1993). We grew strains at permissible (24° C) and non-permissible temperatures (32° C) on YPD and also at the non-permissible temperature with the addition of the microtubule depolymerizing agent, benomyl. The combination of the non-permissive temperature and benomyl led to lethality in the ipl1-2 background (Figure 5A, S10). We next constructed double mutants of ipl1-2 in combination with the outer kinetochore mutations and tested for the rescue of the ipl1-2 phenotype (note these experiments were done a genetic background with yeast histones encoded at the normal chromosomal loci). We observed the tested DASH mutants rescued the temperature and benomyl sensitivities (Figure 5A, S10). We note, ipl1-2 temperature sensitivity was only slightly suppressed by *SPC105*^R583S^ and *NDC80*^E568A^ mutants. These results are consistent with the Nuf2 clustering data that the DASH mutants strongly suppress chromosome instability, whereas the other outer kinetochore complex mutants do not.

**Figure 5.**
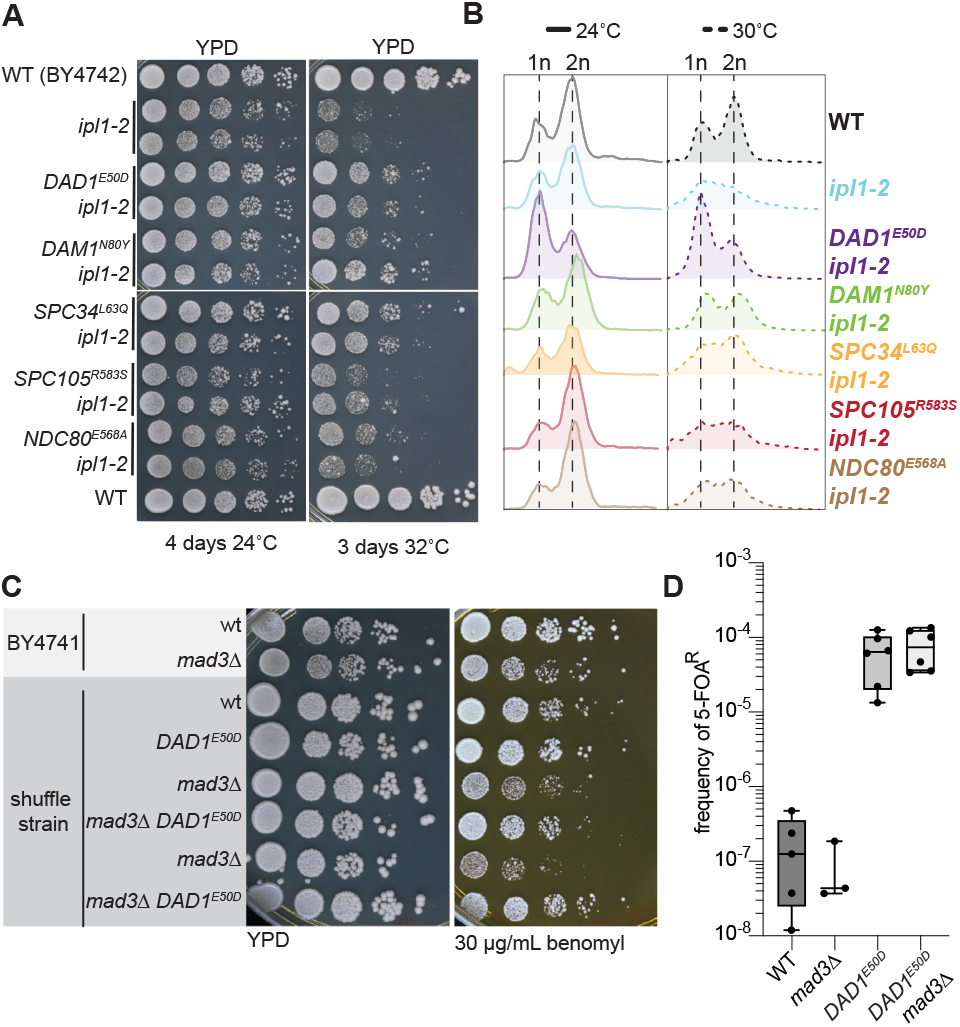
DASH/Dam1c mutants rescue Aurora B kinase and SAC mutants. (**a**) Growth assays of wild type, ipl1-2 mutants, and ipl1-2 kinetochore double mutants at permissive (24° C) and non-permissive temperature (32° C). Tenfold serial dilutions of ~1.0 OD600 yeast cultures were spotted onto YPD solid media and grown at the indicated temperatures for four and three days, respectively. (**b**) Flow cytometry analysis of wild type, ipl1-2 mutants, and ipl1-2 kinetochore double mutants at the permissive (24° C) and non-permissive temperature (30°C). Yeast cultures where grown to logarithmic phase at 24° C and divided and grown at 24° C and 30° C for 6 hours before cells were collected and processed for DNA content analysis using Sytox Green stain. (**c**) Growth assays of wild type, mad3 mutants, and mad3 *DAD1*^E50D^ double mutants on YPD with and without 30 μg/ml benomyl. Ten-fold serial dilutions of ~1.0 OD600 yeast cultures were spotted onto the indicated solid media and grown at 30° C for three days. (**d**) Humanization of mad3 mutant and double mad3 *DAD1*^E50D^ mutant.

Chromosome missegregation in the ipl1-2 strain can be assayed by flow cytometry (Chan and Botstein, 1993). The shift to the non-permissible temperature shows the loss of the distinctive 1n and 2n ploidy peaks of the cycling population (Figure 5B). In agreement with the spot assays, flow cytometry of cells grown at the permissive or non-permissive temperatures showed that the DASH/Dam1c mutants were able to fully rescue the characteristic 1n and 2n peaks of a cycling cell population when grown at 30°C (Figure 5B). However, the *NDC80*^E568A^ and *SPC105*^R583S^ mutants fail to rescue the cycling peaks. We conclude that the DASH/Dam1c mutants not only stabilize ploidy levels outside of the genetic background of histone humanized yeast but also effectively suppress aneuploidy in ipl1-2 cells.

We next investigated whether the DASH/Dam1c mutants could suppress the phenotypic defect of a spindle assembly checkpoint (SAC) mutant, which delays the onset of sister chromosome separation to allow for corrections of unattached or incorrect kinetochore-microtubule connections (Musacchio and Salmon, 2007). We found that the *DAD1*^E50D^ mutation is able to suppress the benomyl sensitivity of the SAC mutant, mad3 (Figure 5C). The humanization of a *DAD1*^E50D^mad3 strain showed that SAC activity was dispensable for robust humanization (Figure 5D). This suggests that the SAC is not active in histone humanized yeasts and that the *DAD1*^E50D^ mutant suppresses chromosome instability independently of this regulatory pathway.

### Molecular basis for suppression of histone-humanization by *DAD1*^E50D^ and *DAM1*^N80Y^

To gain insights into the molecular mechanism of the suppression of chromosome instability, we mapped our mutants onto the cryo-EM structure of *Chaetomium thermophilum’s* DASH/Dam1 complex (Jenni and Harrison, 2018). The mutant residues of Dad1 and Dam1 are separated by less than ~4 Å (Figure 6A), we thus hypothesized that their interaction may underlie a shared molecular mechanism through which they suppress chromosome instability.

**Figure 6.**
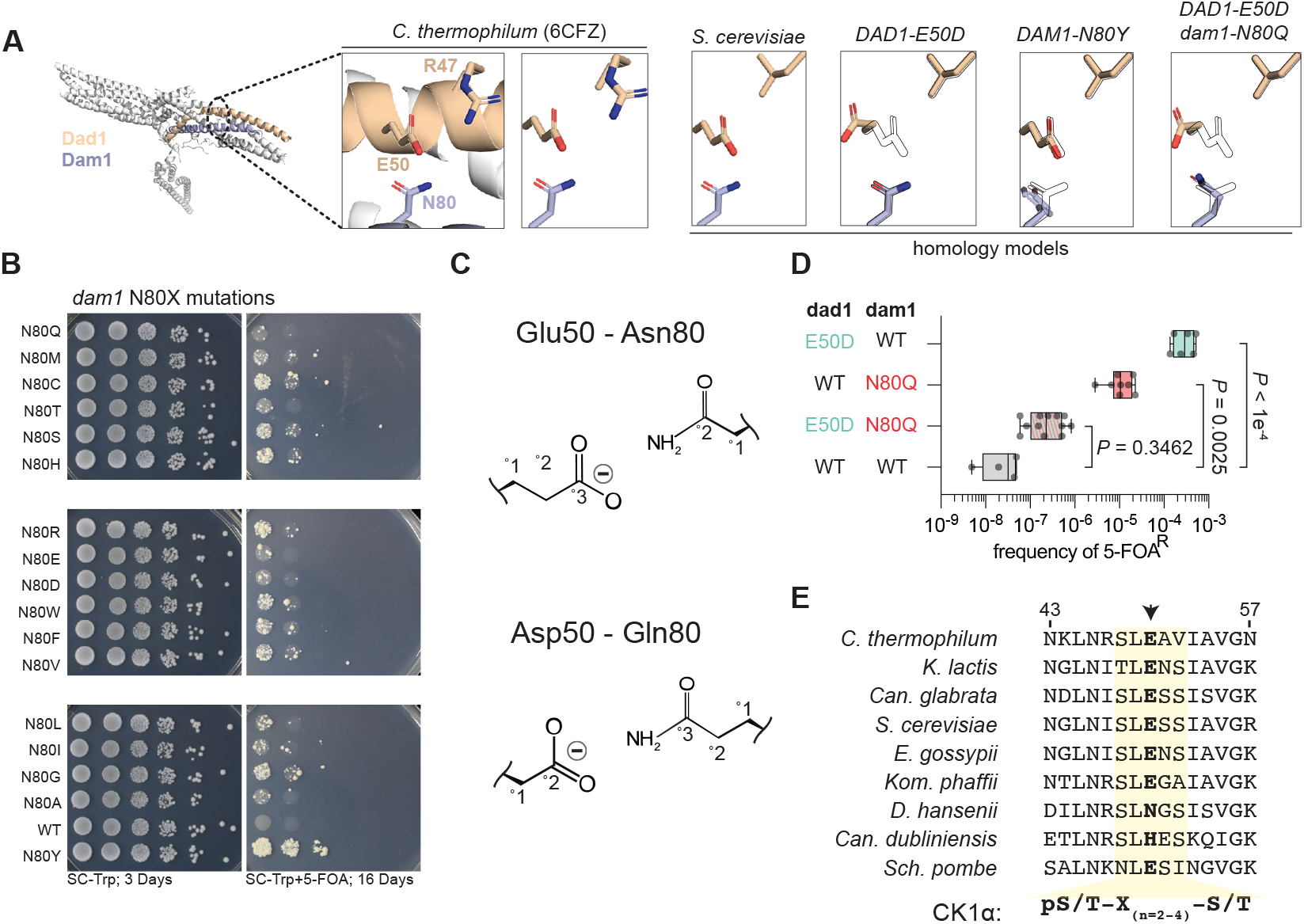
Molecular basis for suppression of histone humanization by *DAD1*^E50D^ and *DAM1*^N80Y^. (**a**) Homology models of *S. cerevisiae* DASH/Dam1c complex. The decametric DASH/Dam1c Cyro-EM structure of C. *thermophilum* is shown, with Dad1 and Dam1 highlighted. The corresponding homology model is shown to the right, with various mutant models shown (with outlined WT residues for reference). Note the relative positioning of Dad1 residue 50 and Dam1 residue 80. (**b**) Histone-humanization assay for various dam1N80X mutants. 5-FOA is used to counter-select the yeast histone plasmid, forcing growth with the human histone plasmid. Yeast were serially diluted from a starting culture of 1.0 OD600. (**c**) Cartoon interactions for complementary pairings of Dad1 residue 50 and Dam1 residue 80. (**d**) The dual mutant Dad1-Asp50 – Dam1-Gln80 DASH/Dam1c fails to humanize, while the single mutants readily humanize. Significance of the mean difference in 5-FOA^R^ frequency was determined with the Mann-Whitney test. (**e**) Protein alignment of Dad1 orthologs, highlighting a conserved casein 1 kinase consensus phosphorylation site (yellow shaded region). Arrow indicates the mutant residue 50 of Dad1.

First, we took a molecular modeling approach to gain insight into the structural changes imparted by the *DAD1*^E50D^ and *DAM1*^N80Y^ mutations. In the cryo-EM structure, the residue Glu50 of Dad1 is stabilized by interactions with the neighboring residue Arg47 (Figure 6A), which is not conserved in *S. cerevisiae* (Figure 6E). We next constructed a homology model of *S. cerevisiae*’s DASH/Dam1c complex to understand how residue Glu50 is arranged in the absence of residue Arg47. In our model, the residue Glu50 of Dad1 interacts exclusively with the residue Asn80 of Dam1 (Figure 6A), potentially via a weak hydrogen bond. Given that either mutation, *DAD1*^E50D^ or *DAM1*^N80Y^, has the potential to disrupt this interaction, we modeled both mutations onto the homology structure of *S. cerevisiae’s* DASH complex. In either case, the mutations resulted in a loss of the WT Glu50-Asn80 interaction (Figure 6A).

Secondly, we experimentally mutated the Asn80 residue of Dam1 to every other amino acid, with the idea being that any mutation at position Asn80 of Dam1 will disrupt the interaction with Glu50 of Dad1. Remarkably, any mutation that we introduced to position 80 of Dam1 significantly improved the rate of humanization over that of WT Dam1, although the original *DAM1*^N80Y^ mutation was superior (Figure 6B S11A). These data support our model that the loss of the Glu50-Asn80 interaction is a key feature for suppression.

We reasoned that we could restore the Glu50-Asn80 hydrogen bond interaction via an alternative amino acid paring, noting that it may be restored by the similar duo of amino acids, Dad1Asp50-Dam1Gln80 (Figure 6C). In agreement, our molecular modeling suggested that the Asp50-Gln80 pairing forms an interaction similar to the WT Glu50-Asn80 interaction (Figure 6A), therefore, the yeast containing both mutations should fail to humanize or humanize at a lower rate than either single mutation alone. As before, we found that the single Dad1-Asp50 or Dam1-Gln80 mutations humanized at rates ~ 10,000-fold and ~370-fold, respectively, over WT (Figure 6D). As predicted, the double Dad1-Asp50 Dam1-Gln80 mutant only weakly humanized at a rate of only ~8.5-fold over WT, significantly worse than either mutation alone, showing that restoration of this hydrogen bond is sufficient to lower the ability to undergo histone humanization (Figure 6D).

However, loss of this hydrogen bond is not the only mechanism at play as it does not account for the difference in humanization rates between the *DAM1*^N80Y^ and *dam1*^N80Q^ variants. We therefore tested if the Dad1 residue 50 glutamic acid was essential for humanization by the *DAM1*^N80Y^ mutation.

To this end, we generated a *dad1*^E50A^ and *DAM1*^N80Y^ shuffle strains and performed the histone humanization assay. As expected the single *dad1*^E50A^ mutation resulted in elevated rate of histone humanization, as predicted due to disruption of the hydrogen bond (Figure S11B). Furthermore, we observed that humanization in the double *dad1*^E50A^ *DAM1*^N80Y^ mutant strain was significantly reduced in comparison to the *DAM1*^N80Y^ alone (Figure S11B). These data suggest that suppression of histone humanization by *DAM1*^N80Y^ functions through additional means that are dependent on Dad1 residue 50 glutamic acid. Taken together, these data point to disruption of the hydrogen bond between Dad1-Glu50 and Dam1-Asn80 being one critical aspect for aneuploidy suppression by the *DAD1*^E50D^ and *DAM1*^N80Y^ mutations.

### DAD1^E50D^ disrupts DASH complex oligomerization and weakens microtubule attachments

What is the mechanism of suppression for the *DAD1*^E50D^ mutation downstream of the loss of the hydrogen bond? We note that the Glu50 residue of Dad1 falls into the middle of an evolutionary conserved region that lies precisely at a critical interface of DASH/Dam1c oligomerization (Figure 6E 7A; Jenni and Harrison (2018)). This suggested that these mutations may alter the oligomerization of the DASH complex. To this end we purified recombinant WT and mutant *DAD1*^E50D^ DASH complexes to characterize the effect of the E50D mutation in vitro. The DASH complex makes visually impressive rings around microtubules in vitro (Miranda et al., 2005) and we wondered if the *DAD1*^E50D^ mutant might perturb this feature. We therefore reconstituted both WT and mutant *DAD1*^E50D^ DASH complexes on microtubules and imaged them using negative stain electron microscopy. We observed that both the WT and mutant *DAD1*^E50D^ DASH complexes formed complete rings around microtubules, but noted that the *DAD1*^E50D^ rings appeared less distinct (Figure 7B). Analysis of single rings confirmed that mutant *DAD1*^E50D^ DASH complex formed rings that are less well defined as indicated by a decrease in the average intensity of pixels near the edges of the microtubule lattice (Figure S12A-B).

**Figure 7.**
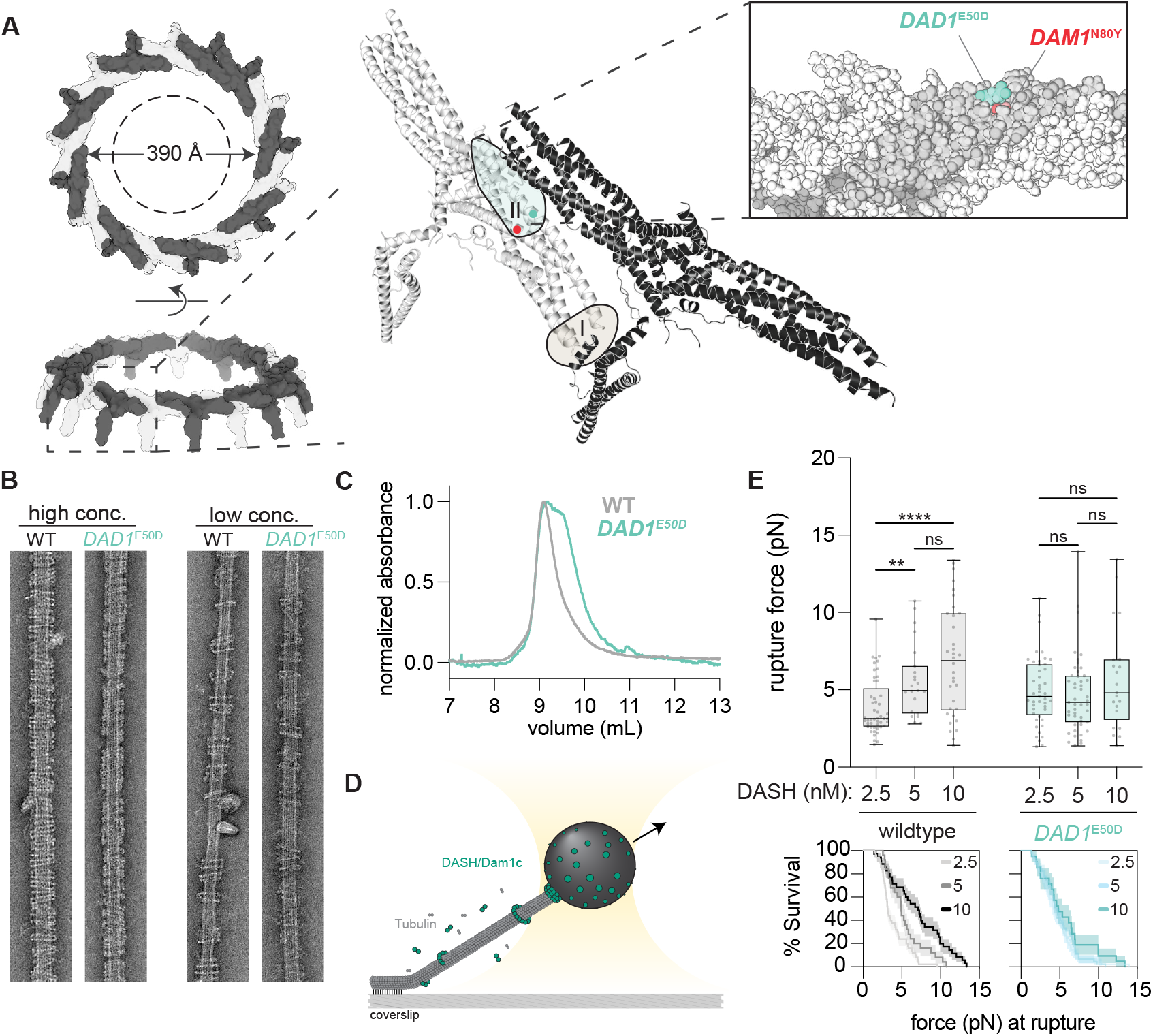
DAD1^E50D^ disrupts DASH complex oligomerization and weakens microtubule attachments. (**a**) Structure of the 17-member DASH/Dam1c ring of *C. thermophilum* is shown (6CFZ). The ring, with a 390 Å interior circumference, encircles a single microtubule with a width of 250 Å (dashed line). Individual protomers oligomerize through interactions of two conserved interfaces, with the DAD1E50 and DAM1N80 residues highlighted in the zoomed in region of interface II. (**b**) Example EM images of DASH rings taken at low and high concentrations. High concentration > 25 nM complex and low concentration 11–16 nM complex. Microtubules are at a concentration of 20 nM tubulin dimer. (**c**) Size exclusion chromatography of the indicated DASH complexes. (**d**) Schematic of the optical trap experiment with wildtype and mutant *DAD1*^E50D^ DASH complex on microtubules. (**e**) Rupture force assay from experiments at three concentrations of DASH complex incubated with the beads. Briefly, a 3.5 pM of beads were incubated with the indicated concentrations of DASH complex (representing a ratio of DASH complex to beads of 714:1, 1428:1, and 2860:1). Below the upper panel are survival curves for each DASH complex are shown at the indicated concentrations.

While the *DAD1*^E50D^ mutant still formed rings, the above single ring analysis suggested oligomerization may be subtly compromised. We hypothesized that the *DAD1*^E50D^ mutant might affect the dimerization of the complex in the absence of microtubules. To test this, we performed size exclusion chromatography for both the WT and mutant *DAD1*^E50D^ complexes. Previous studies have demonstrated that WT DASH complex forms a dimer in solution (Umbreit et al., 2014). Indeed, we observed a single major peak eluting at ~9 mL, corresponding to the WT DASH complex dimer (Figure 7C, S12C). However, for the *DAD1*^E50D^ mutant complex we observed a broadening of this peak with a second major peak appearing at ~9.5 mL, which more clearly appeared at lower concentrations of complex (Figure 7C, S12C). From these data we conclude that the *DAD1*^E50D^ mutation weakens the interaction between DASH/Dam1c protomers, which we observe as an decrease in the dimer fraction and the appearance of a second species of lower apparent molecular weight.

We predicted that the decrease in dimerization ability of the *DAD1*^E50D^ DASH complex would weaken microtubule attachments. We tested the strength of DASH microtubule attachments using a rupture force assay (Figure 7D). We observed for the WT DASH complex that the load bearing strength increased when higher concentrations of complex were added to the beads – in agreement with oligomerization driving stronger attachments (Figure 7E; Umbreit et al. (2014)). However, for the mutant *DAD1*^E50D^ DASH complex we observed no such scaling of strength as all concentrations displayed similar mean rupture forces (Figure 7E). Put together our biochemical and rupture force assays suggest that the *DAD1*^E50D^ mutant weakens the interaction interface between individual protomers, which leads to reduced dimerization and ultimately reduced oligomerization and ring formation to drive weakened microtubule attachments.

### Trade-off between mitotic chromosome segregation versus successful meiosis

We wondered, why if the subtle mutation of *DAD1*^E50D^ improves the fidelity of mitotic chromosome segregation, then why is this interface so well conserved? In particular, why is the Dad1 glutamic acid at residue 50 nearly invariant, if *DAD1*^E50D^ appears more fit in mitosis? To this end we considered meiosis as a potential explanation for why this residue is so well conserved – hypothesizing that the *DAD1*^E50D^ mutation would be detrimental to meiosis. We thus made *DAD1*^E50D^ heterozygote diploid mutant in the sporulation-competent SK1 background and compared the sporulation efficiency to a WT SK1 diploid (Figure 8A). We observed that the *DAD1*^E50D^ mutation severely crippled the efficiency of sporulation, reducing efficiency from ~84% in WT to ~14% in the heterozygous mutant diploid (Figure 8A). In line with reduced meiotic function, we also observed a significant decrease to the viability of spores, observing that the *DAD1*^E50D^ mutation significantly increases the rate of tetrads with fewer than four viable spores (Figure 8B). Intriguingly, in tetrads that produced only two viable spores the frequency of the *DAD1*^E50D^ mutation was significantly enriched over the expected 1:1 ratio to *DAD1*^WT^ (Figure 8C, S13). We conclude that while *DAD1*^E50D^ leads to enhanced fidelity of mitotic chromosome segregation, while it is severely detrimental to the success of meiosis.

**Figure 8.**
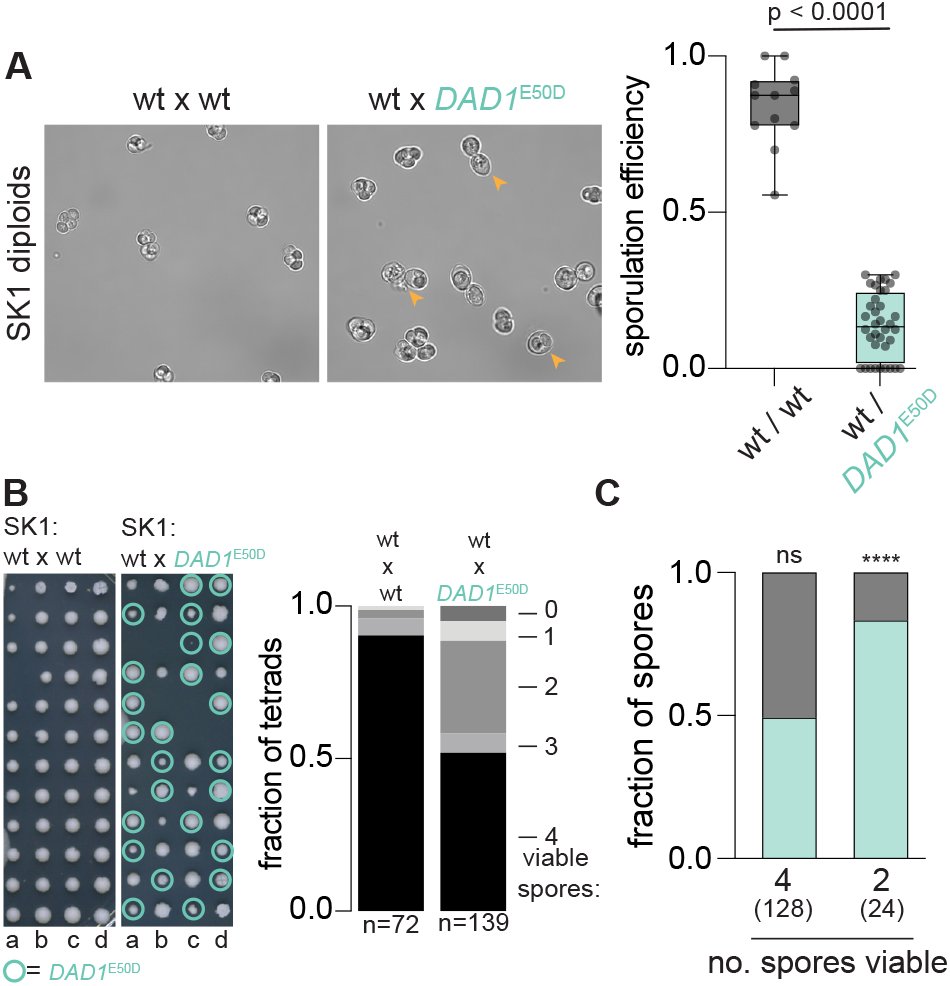
*DAD1*^E50D^ decreases the efficiency of sporulation and spore viability. (**a**) Sporulation efficiency after five days in sporulation medium at 25 ° C. Example micrographs are shown to the left for both wildtype and *DAD1^E50^D* SK1 strains. Right, quantification of average sporulation efficiency from all micrographs. Two tailed unpaired t test of the mean difference in sporulation efficiency. (**b**) Spore viability assay. Individual tetrads were micromanipulated on a agar surface and spores were dissected and arrayed. Genotype was determined by PCR genotyping of the DAD1 locus. Quantification from all tetrad dissections performed, total are shown below. The one-tailed probability of observing 51% spore viability in *DAD1^E50^D* SK1 over a total of 139 tetrads is *p* < 0.000001, assuming a spore viability of ~90% from wild type SK1. (**c**) Genotyping of spores isolated from tetrads with either four viable spores or two viable spores. Fraction of spores with the indicated allele (gray *DAD1*^WT^ and pink *DAD1*^E50D^) is displayed. The one-tailed bionamial test is shown for the probability of observing the frequency of the *DAD1*^E50D^ allele given equal segregation (*DAD1*^WT^, *p* = 0.46; *DAD1^E50D^, p* = 0.0011). Numbers in parenthesis indicates the total number of spores genotyped.

## Discussion

Here we mechanistically determine how the budding yeast *S. cerevisiae* adapts to life with human histones. Following the initial humanization event the genome is shaped by chromosome instability and as a result a high burden of aneuploidy. Selective advantage for any mutation which enhances life with human chromatin is strong, with fitness improving rapidly, but gradually plateauing. We isolate and characterize a class of mutations that arose following the initial humanization event. These mutants increase the fidelity of chromosome segregation in response to centromere dysfunction and ultimately lead to ploidy stabilization and subsequently aneuploidy reduction.

We observed elevated frequency of chromosomal aneuploidy for a non-random set of chromosomes across a meta-analysis of a diverse set of strains. Additionally, we showed that aneuploids are non-adaptive to histone-humanization, suggesting that the frequency of aneuploids is primarily due to some yet-to-be identified centromeric feature(s) that are perturbed by human histones. This unexpectedly led us to the insight that chromosome-specific aneuploid frequency in *S. cerevisiae* may potentially be driven by differences in ancient paralogous centromeres. Mechanistically, it is not clear why this trend between paralog pairs is present. However, it suggests that functional differences in paralogous centromeres have been maintained since the ancient allopolyploidization event that now inform a non-random aneuploid landscape in yeast. Further study will be needed to investigate budding yeast centromere function in the light of centromere paralog evolution.

Successful chromosomal segregation hinges on correct kinetochore–microtubule attachments. As we show, centromere function is disrupted in histone–humanized yeasts – regardless of the specific genetic background examined – yet, the DASH/Dam1c mutants display euploidy. We speculate that in the histone–humanized cells the kinetochore–centromere attachments might be deficient, which could explain a few observations regarding the centromeres of histone–humanized yeasts. First, the centromeres are extremely sensitive to MNase digestion and second, they exhibit a high increase in transcription. Additionally, impaired centromere function helps explain the high rates of chromosome instability in humanized yeast. We note that Mad3, a constituent of the spindle assembly checkpoint (SAC), is totally dispensable for histone humanization, suggesting that the SAC is not active in humanized yeasts. In agreement with the absence of this checkpoint, we observed exceedingly high rates of aneuploidy in histone-humanized lineages, except for the DASH/Dam1c mutants.

How then do the DASH/Dam1c mutants suppress this centromere dysfunction? We propose that the DASH/Dam1c mutants effectively lead to decreased chromosome instability by reducing the strength of microtubule connections. In this model, increased turnover of kinetochore microtubule attachments – induced by weakened DASH complex interactions with microtubules -– allows for reduced errors in segregation (Cimini et al., 2006; Zaytsev and Grishchuk, 2015). This model of correction by detachment, is analogous to how the regulated action of correction by Ipl1 ensures bipolar attachments. Increased cycles of attachment-detachment caused by weakened microtubule-kinetochore coupling may allow cells an opportunity to achieve bipolar attachment. Effectively, this can be thought of as a “blunt” corrective measure, that is potent when the initial rate of biorientation is low or when no mechanism of correction is functioning (i.e., in the absence of SAC/Aurora B activity). Thus, seemingly paradoxically, a compromised inner centromere is effectively rescued by weakening the strength of the outer kinetochore. Further study will reveal deeper insights into how histone–humanized yeast adapt to human histones and the downstream mechanisms of the *DAD1*^E50D^ and *DAM1*^N80Y^ mutants. Of particular interest would be to assess whether kinetochores from histone-humanized yeasts can sense tension or whether Aurora B is inactive in histone-humanized yeasts. Our data is consistent with Aurora B being inactive histone-humanized yeasts as we observed high rates of chromosome aneuploidy and that our DASH/Dam1c mutants suppress the loss of Aurora B activity (ipl1-2). We speculate that the intrinsic catch bond-like behavior of yeast kinetochores may remain functional in histone-humanized yeasts, which when coupled alongside *DAD1*^E50D^ induced kinetochore-microtubule turnover, may allow for stabilization of proper kinetochore-microtubule attachments (Akiyoshi et al., 2010).

The phenotypic effects of the subtle *DAD1*^E50D^ mutation, from a glutamic acid to aspartic acid, are dramatic. It underlies a potent adaptive route to life with human histones – which we propose is due to suppression of centromere dysfunction. Furthermore, *DAD1*^E50D^ mutation is not only limited to the peculiarity of the histone–humanized genetic system, as it suppresses chromosome instability and aneuploidy accumulation in genetic backgrounds with yeast histones (Figure 5). Our data collectively supports the notion that *DAD1*^E50D^ mutation enhances the fidelity of chromosome segregation during mitosis, especially during periods of chromosome instability, however, it is crippling to the success of meiosis. Interestingly, the *DAD1*^E50D^ mutation lies within a conserved predicted phosphorylation site of the casein kinase I Hrr25, which is a component of the meiosis I-specific kinetochore crosslinking monopolin complex (Figure 6E; Sarangapani et al. (2014)). Whether or not Dad1 is a target of Hrr25 or if the *DAD1*^E50D^ mutation interferes with Hrr25 meiotic function is not known. However, our observation that *DAD1*^E50D^ disrupts meiosis suggests that this interface maybe critical for successful meiosis and it will be of interest to investigate if DASH oligomerization is regulated during the transition from meiosis I kinetochores to meiosis II ones.

During preparation of our manuscript Clarke et al. reported strikingly similar conclusions from a completely orthogonal genetic screen for elevated rates of aneuploidy (Clarke et al., 2022). Notably, they developed a screen for suppressors of deficiency in BIR1, encoding a component of the chromosomal passenger complex, deletion of which leads to increased chromosome segregation defects and elevated aneuploidy. Remarkably, they isolated a distinct set of DASH complex mutants targeting the same oligomerization interface. Furthermore, they described a second pathway of bir1-induced chromosome instability suppression through mutation of the chromosomal passenger complex (CPC) subunit Sli15 (G334S), which increases its microtubule binding independently of its centromere recruitment. We observed that in our loss of aneuploidy experiment the *DAD1*^E50D^ mutant was lost and supplanted by the Sli15 (D331Y) mutant, perhaps due to direct recruitment of the CPC to microtubules alleviating the benefit of the *DAD1*^E50D^ mutation. Our results complement and extend their findings to reveal the molecular and biochemical details demonstrating that weakened kinetochore–microtubule attachments provides a mechanistic path for suppression of dysfunctional centromeres. In particular, our in vitro data directly illustrates that the *DAD1*^E50D^ mutation drives weakened microtubule attachments via impairment to DASH/Dam1c oligomerization.

In sum, our data reveal a mechanism for suppression of human histones through weakened microtubule connections in budding yeasts. We note that this is only one of many paths through which yeast may adapt to human histones, as we identified multiple cellular processes which are enriched in our list of candidate suppressor mutations. (Figure S2; Supplemental table 1). Lastly, we highlight the unique evolutionary forces at play in histone humanized yeasts which not only illuminate the centrality of nucleosomes in eukaryotic life but afford a powerful system in which to probe biological functions.

## Methods

### Strains, plasmids and oligos

All strains used in this work are listed in supplementary table 6 and are available upon request. To construct the haploid and diploid shuffle strains with human histone suppressor mutations we used CRISPR-Cas9 to scarlessly introduce the mutation of interest. First, we designed sgRNAs targeting our genes of interest alongside repair templates to introduce the desired mutations (Figure S3A). To clone the guide RNAs we designed a single oligo that consisted of 20 bp upstream the nearest NGG to our codon of interest, flanked by homology sequences for Gibson assembly into our expression plasmid (5’–TGAAAGATAAATGATC–20bp–GTTTTAGAGCTAGAAA). For the Gibson assemblies, 1 μg of plasmid DNA was digested with NotI and CIP overnight and then purified with the Zymo DNA Clean and Concentrator kit (Zymo Research cat. D4030). Next, to the Gibson mix we added 20 ng of digested plasmid with 1 uL of the sgRNA oligo (50 μM), mixed and incubated at 50°C for 1 hour. The Gibson reaction was then diluted 1:10 and the dilution was transformed into chemically competent E. coli, the transformation was plated to the appropriate selection and clones verified by sanger sequencing.

We used a double stranded DNA (dsDNA) repair template to introduce the mutation and eliminate the PAM site. dsDNA was constructed from two oligos that are annealed and 5’ overhangs filled in using Klenow polymerase according the manufacture’s specifications (NEB cat. M0210L). All yeast transformation were carried out using the lithium-acetate method. For each CRISPR edit, the strain of interest was first pre-transformed with a Cas9 plasmid, and cells harboring Cas9 were transformed with the guide RNA plasmid and the dsDNA repair template. Edited cells were then selected by double selection of the Cas9 plasmid and the sgRNA plasmid. Strains were validated by sanger sequencing.

All plasmids are listed in supplemental table 7. All oligos are listed in supplemental table 8.

### Humanization assay and evolution experiments

We humanized yeast’s core histone genes as previously reported (Haase et al., 2019; Truong and Boeke, 2017). Briefly, a shuffle strain – which has the four histone gene clusters chromosomally deleted and has a single set of yeast histone genes on a *URA3* plasmid – is transformed with a *TRP1* plasmid encoding a single set of human histone genes. Importantly, the yeasts and human histone genes are expressed using orthogonal promoters and terminators. Next, each transformant is inoculated into 5 mL of SC-Trp medium containing 2% galactose and 2% raffinose as the carbon source, this is to inactivate the conditional centromere on the yeast-histone *URA3* plasmid.

Once the culture has reached saturation, a range of volumes (typically 1uL for a robust humanizing strain or up to 1mL for a weak humanizing strain) are plated to SC-Trp agar plates containing 5-FOA (1 mg mL-1), to counter select the *URA3* marker (Haase et al., 2019). Plates are sealed in a Tupperware box with a damp paper towel and growth at 30°C is monitored for up to 4 weeks, but longer in some cases. After a final count of colonies which grew on the SC-Trp+5-FOA plates a humanization rate is calculated – the total number of colonies is divided by the estimated number of cells plated.

Colonies which appeared after 7 days of growth on plates with 5-FOA were PCR genotyped to confirm the loss the yeast histone genes (Haase et al., 2019). Confirmed clones were patched to rich media agar plates (YPD) and stored in glycerol at −80°C. A portion of each patch was then cultured in SC-Trp and grown to saturation (generation 0). Cultures were then diluted by inoculated 5 mL of growth medium with 50 μL of saturated culture, this passaging was repeated for 5 cycles. At each time point a portion of the culture was stored in glycerol at −80°C.

### Whole genome sequencing and variant analysis

Genomic DNA was extracted from stationary yeast cultures using a double phenol-chloroform extraction method. Briefly, yeast pellets, from ~1.5 – 5 mL of culture, were disrupted by bead beating in a solution consisting of 225 μL 1x TES (TE buffer + 0.5% SDS) plus 200 μL of 25:24:1 phenol:chloroform:isoamyl alcohol (Thermo Fisher cat. 15593031) in tubes with a pre-aliquoted amount of 0.5 mm diameter yttria-stabilized zirconium oxide beads (MP Bio cat. 116960050-CF). A second step of phenol-chloroform extraction was done for each sample and DNA was precipitated in 70% ethanol. Extracted genomic DNA (gDNA) was then resuspended in a solution containing 30 mg/mL RNAse A (Thermo Fisher cat. EN0531) to remove any RNA. Next, approximately 50 ng of gDNA was used as input to the NEB Ultra II FS Library Prep Kit for Illumina (NEB cat. E7805L). Libraries were then sequenced using either paired end 2×36 bp or 2×72 bp read chemistry. Single nucleotide variants were called as previously described (Truong and Boeke, 2017).

### Ploidy estimate from whole genome sequencing data

Chromosomal ploidy was estimated from the sequencing coverage data. First, the median coverage of 1 kb windows across the genome was calculated. From these windows, we divided each by the median coverage of the euploid genome (those chromosomes for which we never observed aneuploidy in humanized strains: VII, XIV, XIII, and XV) and took the log2 ratio. The mean of these log2 ratio windows for each chromosomes was used to determine ploidy counts. In some cases, especially for lower sequencing coverage samples we observed a smiley pattern of coverage (Gallone et al., 2016), in such situations we had to manually annotate ploidy levels.

### Growth assays

Because the humanized strains displayed rapid improvement in growth rates and variability across the population, we avoided measurements of growth in liquid culture to ensure that a “rare” fast grower would not take over the population. We therefore measured growth rate as the rate of change in the area of colonies grown on a YPD agar surface. To ensure accurate growth rate measurements of each generation, no preceding step of culturing was performed prior to measurement. To this end we struck out each generation from −80°C glycerol stocks to YPD agar plates. Scans were acquired using the ScanMaker 9800XL Plus (Microtek International) for each generation at two time points to calculate the rate of change in mm^2^ hr^-1^. Colony sizes were measured in the image analysis program Fiji (Schindelin et al., 2012), by manual analysis.

### Ploidy assessment by flow cytometry analysis

Cultures were grown to mid-log phase and 1 – 2 x 10^7^ cells were collected by centrifugation and resuspend in 1.5 mL of water. Next, in order to fix and permeabilize the cells 3.5 mL of 100% ethanol was added slowly to each and left overnight at −20°C. Then, cells were pelleted and washed 3 times with water. To remove contaminating RNA, suspensions were centrifuged and resuspended in a RNAse A solution (15 mM NaCL, 10 mM Tris pH 8.0, 0.1 mg/mL RNAse A) and incubated at 37°C overnight. Cells were then pelleted and resuspend in 50 mM Tris. 0.5 mL of processed cells were then mixed with 0.5 mL of SYTOX Green stain (2 μM SYTOX Green (Thermo Fisher cat. S7020) in 50 mM Tris pH 7.5) and incubated for one hour at 4°C in the dark. Cells were finally pelleted and resuspended in 1 mL of Tris pH 7.5 and sonicated. Flow cytometry analysis was performed on the BD Accuri C6 flow cytometer and the data was analyzed in the program FlowJo (v10.0.7).

### RNA extraction, total RNA sequencing, and analysis

Yeast strains were grown at 30°C to mid log phase (between OD600 0.6 - 0.8) in biological triplicate. Cultures were immediately placed on ice, pelleted, and flash frozen in liquid nitrogen then stored at −80°C. Total RNA was then extracted by resuspend each cell pellet in RNA extraction buffer (50 mM Tris-HCL pH 8.0 and 100 mM NaCL) and transferred to a 2 mL screw capped tube containing 0.5 mm diameter yttria-stabilized zirconium oxide beads and samples were mechanically disrupted at 4°C for 15 cycles of 5 seconds shaking and 30 seconds rest. DNA and RNA was then purified by phenol chloroform extraction, by mixing at a 1:0.95 ratio of lysate to a 25:24:1 mixture of phenol:chloroform:isoamyl alcohol. To precipitate the nucleic acid the aqueous fraction was then mixed with 99.5% ethanol to a final solution of 70% ethanol. Purified DNA and RNA was then treated with DNAse I (Agilent cat. 600031) to remove contaminating DNA from each sample. RNA quality was finally assessed on an agarose gel.

Purified RNA was then used for preparing total-RNA stranded RNAseq libraries with the QIAseq Stranded Total RNA Lib Kit (Qiagen cat. 180745) according to the manufacture’s specification. Ribosomal rRNA was depleted using the QIAseq FastSelect –rRNA Yeast Kit (Qiagen cat. 334217). Libraries were sequenced as paired-end 75 base pair reads on a Illumina NextSeq 500. Reads were first processed to remove Illumina barcodes and quality trimming with Trimmomatic (v0.39) and quality assessed with FastQC (v0.11.4). Processed reads were then aligned to the S288C reference genome (release R64-2-1) using the Kallisto pseudoalignment (v0.46.0) for CEN RNA transcript counts and STAR aligner (v 2.5.2) for *CEN* RNA genome tracks.

### MNase sequencing analysis

MNase digested raw sequencing data was downloaded from our previous study and reads were processed as before (Truong and Boeke, 2017). Aligned reads were then used as input into the DANPOS program (Chen et al., 2013) to estimate nucleosome occupancy and positioning across the genome. Finally, profile tracks for the sixteen centromeres were made using the profile function and results were plotted using MATLAB. Orientation of all centromeres when plotted was kept to run from CDEI to CDEIII.

### *CEN* RNA RT-PCR analysis

Total RNA was extracted as before from log phase cultures. cDNA was generated using the SuperScript™ IV Reverse Transcriptase (Thermo Fisher cat. 18090010) according to the manufacture’s specifications using a polydT20 oligo in order to enrich for polyadenylated transcripts including *CEN* RNAs. Reversed transcribed RNA was then used for PCR amplification using *CEN* RNA specific oligos and GoTaq® Master Mix. Amplification was performed as follows; 20 cycles of 98°C – 15 sec, 55°C – 15 sec, and 72°C – 15 sec. DNA was analyzed on 1% agarose TTE gel and image was acquired on a BioRad GelDoc.

### RFP-Nuf2 strain construction and imaging

To image kinetochores and the spindle pole body we scarlessly tagged Nuf2 and Spc110 with RFP and GFP, respectively. Briefly, each histone shuffle strain, pre-transformed with a plasmid encoding Cas9, was transformed with a guide RNA expressing plasmid targeting the 5’ end of the gene and a circular dDNA repair template that encoded the appropriate fluorescent protein. The repair templates encoded a fluorescent protein gene (ymScarlet or mNeonGreen) and homology sequences targeting 500 bp up and downstream from the stop codon. Correct repair of the double strand break results in C-terminal fusion genes and strains were validated by imaging and PCR genotyping. Imaging was done in asynchronous populations grown to mid-log phase (OD600 0.6 - 0.8). Histone humanized strains were generated as above.

### DASH complex homology modeling

Homology models were constructed in the SWISS-MODEL suite (Waterhouse et al., 2018). To obtain models for the various Dad1 and Dam1 mutants we chose to model only the arm II of the complex (consisting of the proteins Dad11–94, Dam162–115, Duo159–121, Dad36–94, and Spc345–38) in a 1-1-1-1-1 hetero-state, using the know structure 6cfz.1 as the template. For each mutant model considered, we did the same with the only difference being that the query peptide of interest was modified to contain the mutant residue.

### Dam1 residue 80 CRISPR/cas9 editing

In order to generate mutations at residue 80 on Dam1 (N80X), we used CRISPR/cas9 genome editing as above. Briefly, a guide RNA targeting Dam1 was used to introduce a double strand break, which we repaired with dsDNA donor templates encoding both the mutation of interest and a PAM inactivating silent mutation. We were able to isolate 16 out of 18 possible mutations (excluding the wild type asparagine and the already cloned tyrosine variant). For two mutations, N80P and N80K, we only ever isolated clones that inactivated the PAM sequence but retained the wild type residue 80 sequence – we did not pursue these further.

### DASH complex purification and tension measurement

*Saccharomyces cerevisiae* DASH complex was expressed in Escherichia coli using a single polycistronic vector, as previously described (Flores et al., 2022). The Spc34p component of the DASH/Dam1 complex contained either a C-terminal FLAG-tag or a His6-tag. BL21 Rosetta 2 DE3 cells were transformed with either wild-type DASH complex (pJT44 – FLAG, or PC4 – 6X His) or with single mutations described in the manuscript. Cells were grown to OD600 of 0.6, where the addition of 2 mM IPTG induced protein expression. Cells were induced for 16 hrs at 18°C while shaking at 240 rpm. Cells were collected by centrifugation, resuspended in 50 mM sodium phosphate buffer (pH 6.9) containing 500 mM NaCl, 1 mM PMSF, and 1 protease tablet (Roche) and lysed with a French press. Lysates were cleared by centrifugation (20 min, 25,000 x g, 4°C). Mutant and wild-type DASH complexes were first purified via affinity chromatography using either FLAG resin (GE Healthcare Bioscience) or Nickel resin (Bio-Rad). For FLAG purification, lysate was applied onto FLAG resin under gravity flow and eluted with 50 mM sodium phosphate buffer (pH 6.9) containing 500 mM NaCl and 100 μg/mL FLAG peptide. For His-tagged DASH complexes, samples were eluted in 50 mM sodium phosphate buffer containing 500 mM NaCl and 500 mM imidazole. Both His and FLAG tagged DASH constructs were further purified via size exclusion chromatography.

Recombinant His6-tag DASH complex was linked to 0.56 μm-diameter streptavidin-coated polystyrene microbeads (Spherotech) using biotinylated His5-antibody (Qiagen), as previously described (Asbury et al., 2006; Franck et al., 2007; Powers et al., 2009). For rupture force assays, a final concentration of 2.5, 5, or 10 nM DASH complex was incubated with 3.5 pM beads for 1 hour. Beads were washed twice with AB solution (1X BRB80 [80mM potassium PIPES buffer, pH 6.8, containing 1 mM MgCl2, and 1 mM EGTA], 2 mg/mL BSA, and 1 mM DTT) to remove unbound His6-tagged DASH complex. Glass slides and functionalized coverslips were used to construct flow channels. Channels were functionalized by adding 5 mg/mL biotinylated BSA (Vector Labs) and incubated for 15 min inside a humidity chamber, washed with 1X BRB80, and then incubated with 0.3 mg/mL avidin DN (Vector Labs) for 5 min. Following, channels were further washed with 1X BRB80, and biotinylated microtubule seeds stabilized with GMPCPP were added and allowed to incubate for 5 min inside the humidity chamber. Growth buffer (1X BRB80, 8 mg/mL BSA, 1 mM GTP, and 1 mg/mL -casein) was added and allowed to incubate in the chamber for 5min. DASH-coated beads were added to a reaction mix (1X BRB80, 8 mg/mL BSA, 1 mM GTP, 40 mM glucose, and 1 mM DTT) and sonicated for 10 s. Following sonication, an oxygen scavenging system (250 μg/mL glucose oxidase, 30 μg/mL catalase, and 4.5 mg/mL glucose) was added. Lastly, ~2-2.5 mg/mL of purified bovine brain tubulin was added to the reaction mixture as well as 5 μM FLAG-tagged DASH complex. Reaction mixture was subsequently added to the channel, which was sealed with nail polish.

Data collection was performed as previously described (Flores et al., 2022). Briefly, data was collected for a total of 1 h after the addition of tubulin into the reaction mixture. To determine whether a DASH complex-coated bead was able to bind to microtubules in the absence of force, it was placed onto a microtubule and then trap was shuttered. Microtubule-attached beads were initially pulled at 1 pN of force. After, a gradual increasing force was applied (0.25 pN/s) until the bead ruptured from the microtubule tip or the maximum trapping force was reached (~20 pN under the conditions described here). Rupture forces were analyzed using Igor Pro (WaveMetrics).

### Electron Microscopy

EM was performed as described below, based on the methods outlined in Kim et al. (2017). Bovine brain tubulin was clarified at a concentration of 75 μM at 4°C in BRB80 (80 mM potassium PIPES buffer (pH 6.8), 1 mM EGTA, 1 mM MgCl2) at 90,000 rpm in a Beckman TLA-100 rotor for 10 minutes. From the supernatant, 26.5 μL of clarified tubulin was removed and 1.5 μL 100 mM MgCl2, 1.5 μL DMSO plus 0.5 μL GTP was added. The resulting solution was incubated at 37° C for 30 minutes to allow polymerization of tubulin. A wide bore pipette was used to add 140 μL of BRB80 containing 10 μM taxol (BRB80-T) to the resulting microtubules. The reaction was mixed by gentle pipetting and 100 μL was spun at 58,000 rpm at 37° C in a Beckman TLA-100 rotor for 10 minutes to pellet the microtubules. Supernatant was removed and microtubules were resuspended in 100 μL BRB80-T using a wide bore pipette at room temperature. MT concentration was calculated via BCA assay of the resuspended MT pellet.

Samples for EM were prepared in a final volume of 50 μL BRB80-T containing 20 nM microtubules plus between 5 and 35 nM DASH complex (WT or E50D). Once mixed, MTs plus DASH complex were incubated at room temperature for 30 minutes. Carbon-coated copper grids were negatively discharged in a glow discharge device. A 3 μL volume of sample was applied onto a discharged grid for 30 s before excess liquid was blotted away with Whatman 1 filter papers. The grid was then washed with BRB80-T and blotted three times prior to application and blotting of 30 μL of 2% uranyl formate three times. The stained grid was then air dried prior to imaging.

The grids were imaged on a Tecnai T12 transmission electron microscope (FEI, Hillsboro, OR) operating at 120 kV using a Tungsten filament electron source. Images were recorded on a bottom-mounted Ultrascan 4000 (Gatan, Pleasanton, CA) camera at 42,000x nominal magnification. The Fiji implementation of ImageJ was used to export images in PNG format for publication.

Image analysis of oligomerized rings was carried out in Fiji using the plot profile function to obtain the intensity profiles along the distance of individual DASH complex rings (with the midpoint set at the center of the microtubule lattice). Intensity data was than normalized by subtraction of the average intensity divided by the standard deviation of intensity. The normalized average intensity profile of the unbound microtubules was then subtracted from each to obtain the profiles corresponding to the DASH complex rings.

### DASH complex Size Exclusion Chromatography

Size-exclusion chromatography was carried out using a Superdex 200 Increase 10/300 GL column (Cytiva) equilibrated in DASH complex purification buffer (500mM NaCl, 50mM sodium phosphate buffer (pH 6.9). The column was calibrated using a gel filtration markers kit for protein molecular weights 29,000-700,000 Da (Sigma-Aldrich, product number MWGF1000).

### Sporulation and ascus dissections

Diploid yeasts were grown for two days on solid GNA agar plates prior to sporulation induction. Yeast were then resuspended to 1 x 10^7^ cells in 5 mL of sporulation medium (1% potassium acetate, 0.05% zinc acetate, with supplemented amino acids) and rotated for 5 days and 25°C. After 5 days, cultures were either immediately processed or stored at 4°C for up to two weeks. Asci were digested in 25 μL of 0.5 mg/mL zymolase-20T in 1M sorbitol for 6–7 minutes at 37° C, at which point 200 μL of 1M sorbitol was gently added and digested asci placed on ice. Spores were then micromanipulated and arrayed onto YPD agar plates.

## Data deposition and availability

Raw sequencing reads are deposited and available at the Sequence Read Archive (SRA – Bioproject accession number: PRJNA884379 and Accession numbers: SRR21712309 - SRR21712362).

## Acknowledgements

We thank Imani Sinclair and the NYU Reagent Preparation core for their support with growth media used in this study; Joel Quispe for help with EM and his maintenance of the UW Beckman CryoEM Center; Hannah Ashe, Gwen Ellis, Raven Luther, and Matthew Maurano for providing Illumina sequencing service. This work was supported by the National Science Foundation (URoL-1921641) to J.D.B, NIGMS R35130293 to T.N.D. and NIGMS R35134842 to C.L.A.

## Contributions

M.A.B.H conceptualization, design, experimental work, formal analysis, figure preparation, original manuscript draft, editing. R.L.F. experimental work, formal analysis, editing. G.O. experimental work, formal analysis, editing. E.B-A experimental work, formal analysis. A.Z. experimental work, formal analysis, editing. M.S.D. experimental work. L.L.S experimental work, formal analysis, editing. C.L.A design, supervision, funding, editing. T.N.D. design, supervision, funding, editing. D.M.T experimental work, funding, editing. J.D.B conceptualization, design, funding, editing.

## Supplementary Information

**Figure S1.**
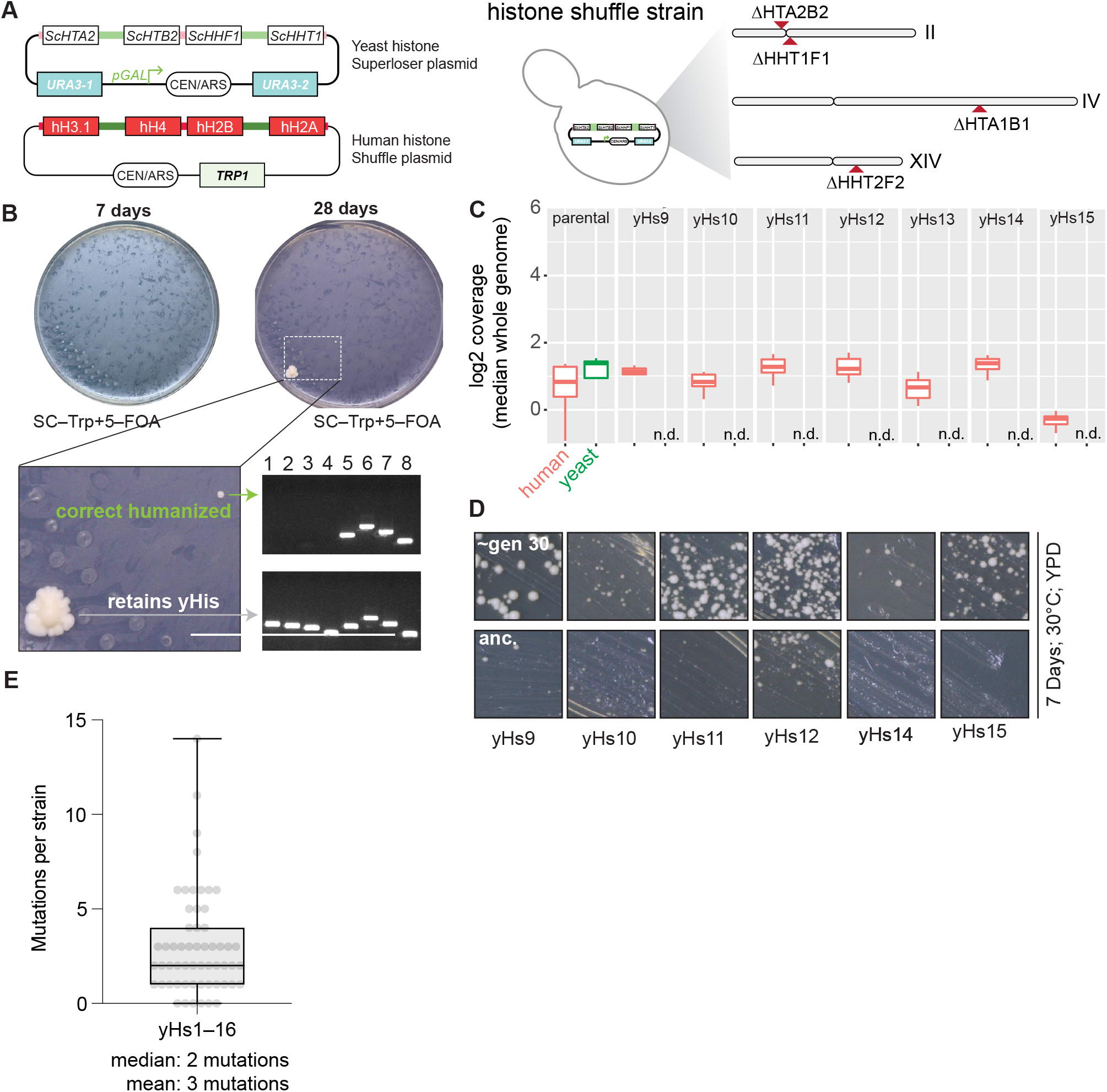
Isolation and verification of histone-humanized yeasts. (**a**) Dual-plasmid histone shuffle assay overview. First a strain with all core histone gene clusters deleted is maintained with a single set of yeast histone genes on the counter-selectable *URA3* plasmid (Haase et al., 2019). In order to shuffle out the yeast histones for human histones, a second plasmid encoding the four core human histones and with a *TRP1* selectable marker is transformed into the shuffle strain. The shuffle strains in then grown on media containing 5–FOA to force the cells to grow with exclusively human histones. (**b**) Example humanization experiment results. Plates are shown at two time points to illustrate the severe growth defects upon the initial humanization event. Two colonies were isolated, one retaining all yeast histones (either through inactivation of *URA3* or some plasmid recombination event) and a bona fide histone–humanized clone as verified by PCR genotyping (Haase et al., 2019). (**c**) Whole genome sequencing coverage plots of the two histone plasmids (human histones, red; yeast histones, green) are shown for the parental shuffle strain prior to humanization and for histone–humanized clones (yHs9–yHs15). (**d**) YPD plate growth assays for histone–humanized clones from the original unevolved glycerol stock and the evolved descendants after ~30 generations of growth. (**e**) Box and whisker plots of number of mutations observed per histone–humanized lineages.

**Figure S2.**
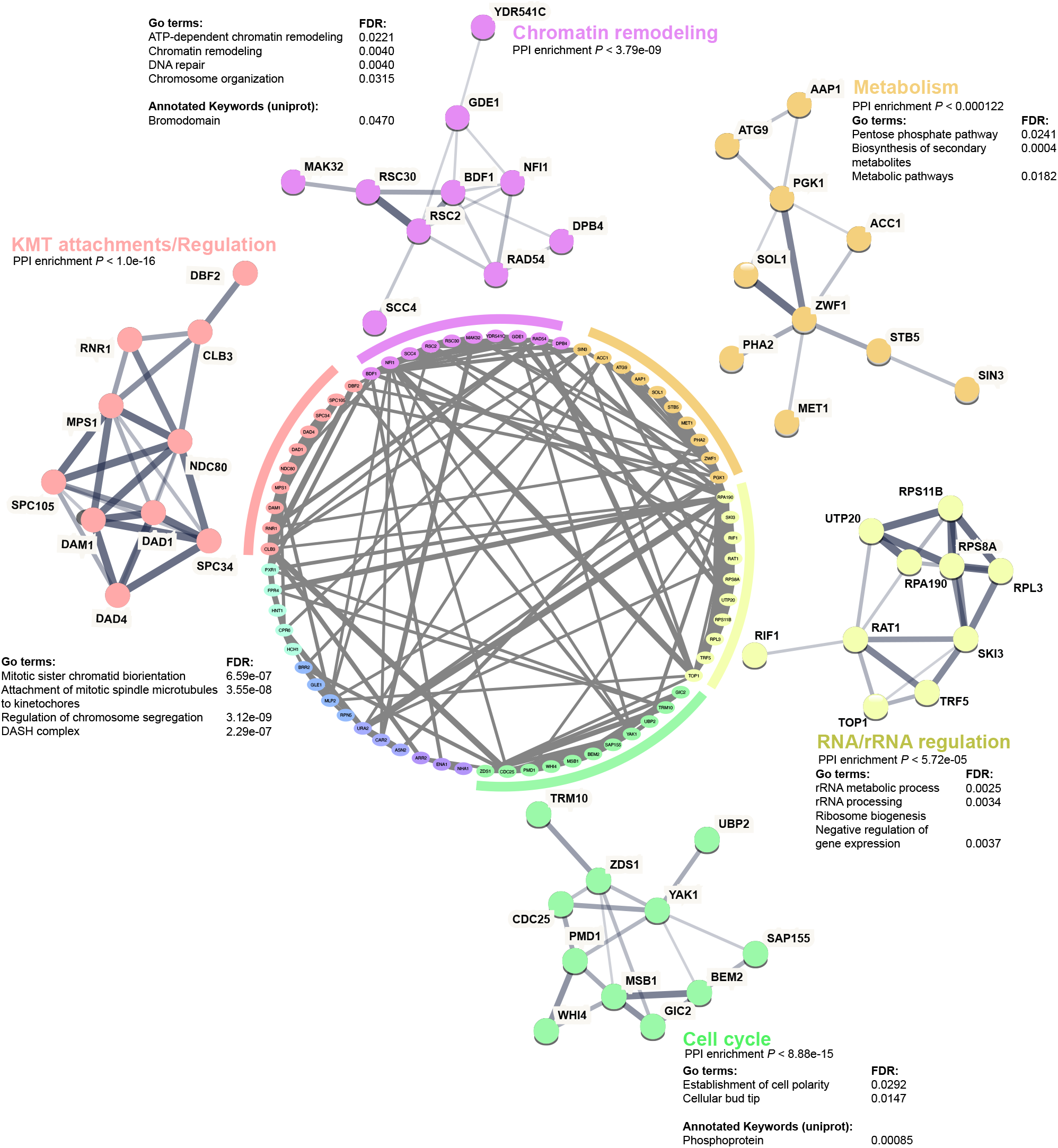
Protein-protein interaction network of candidate suppressors of human histones. (**a**) String database analysis of protein-protein interactions (PPI) considering missense, nonsense, and noncoding mutations identified in histone humanized lineages (Supplemental table 1 and Truong and Boeke (2017)). The network was clustered using MCL clustering with the inflation parameter set to 1.8, clusters are colored, with PPI depicted as gray connecting lines. The number of PPI in the global network was significantly enriched from background (PPI enrichment *p* = 0.0049). Local PPI networks are shown for clusters containing more than 10 interactors, with PPI enrichment p-values shown, alongside Go term enrichment terms and Uniport keywords. Network was constructed using the interaction sources: “textmining”, “experiments”, “databases”, and “co-expression”. The width of each edge represents the strength of interaction between nodes based on available evidence. Disconnected nodes were removed for clarity.

**Figure S3.**
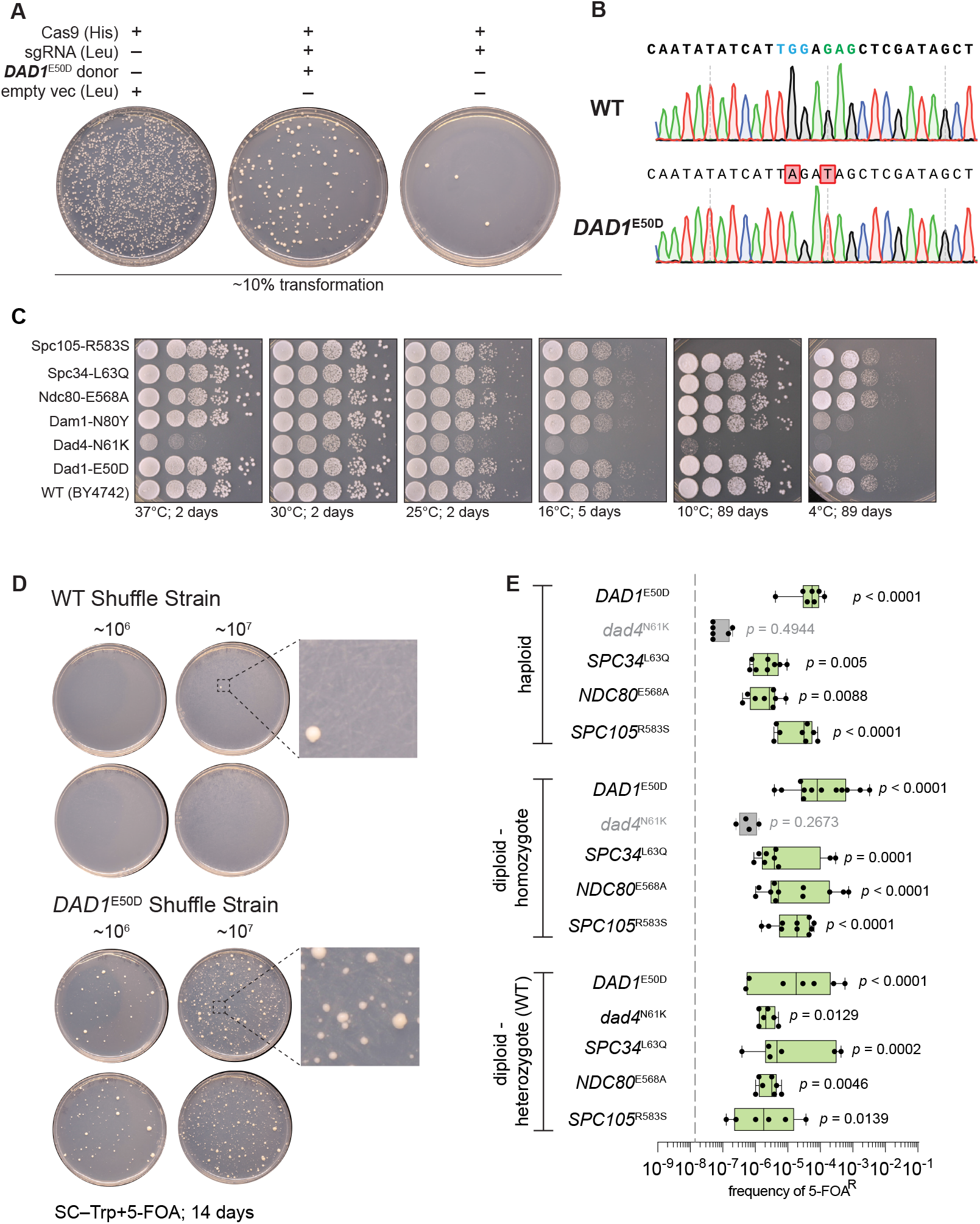
CRISPR-Cas9 mediated mutation of outer kinetochore genes and suppression validation. (**a**) Example CRISPR/Cas9 editing transformation to scarlessly introduce each point missense mutation to an isogenic histone shuffle strain. Note in the absence of dsDNA donor the sgRNA targeting *DAD1* results in a severe killing phenotype and upon co-transformation with a proper dsDNA the killing phenotype is rescued. (**b**) Example Sanger tracks for the edited *DAD1*^E50D^ mutation and wild type sequence. The targeting PAM is colored blue and edited codon in green. (**c**) Growth assay on rich medium (YPD) for the indicated histone shuffle strains with yeast histones. Spots are 10-fold serial dilutions from starting OD600 of 1.0. (**d**) Example suppressor mutation sufficiency histone-humanization experiment is shown for the *DAD1*^E50D^ mutation. Two concentrations of cells were plated (10^6^ mL^-1^ and 10^7^ mL^-1^). (**e**) Histone–humanizations for all tested suppressor mutations are shown. The average rates for wild type strains are plotted as a gray dashed line. Significance was determine with a Kruskal-Wallis test of the mean frequency of 5-FOA^R^ for each mutant versus the mean frequency of 5-FOA^R^ of wild type, with multiple comparisons corrections with the false discovery rate method. Green colored boxes represent suppressors who significantly increased the rate of humanization above wild type level.

**Figure S4.**
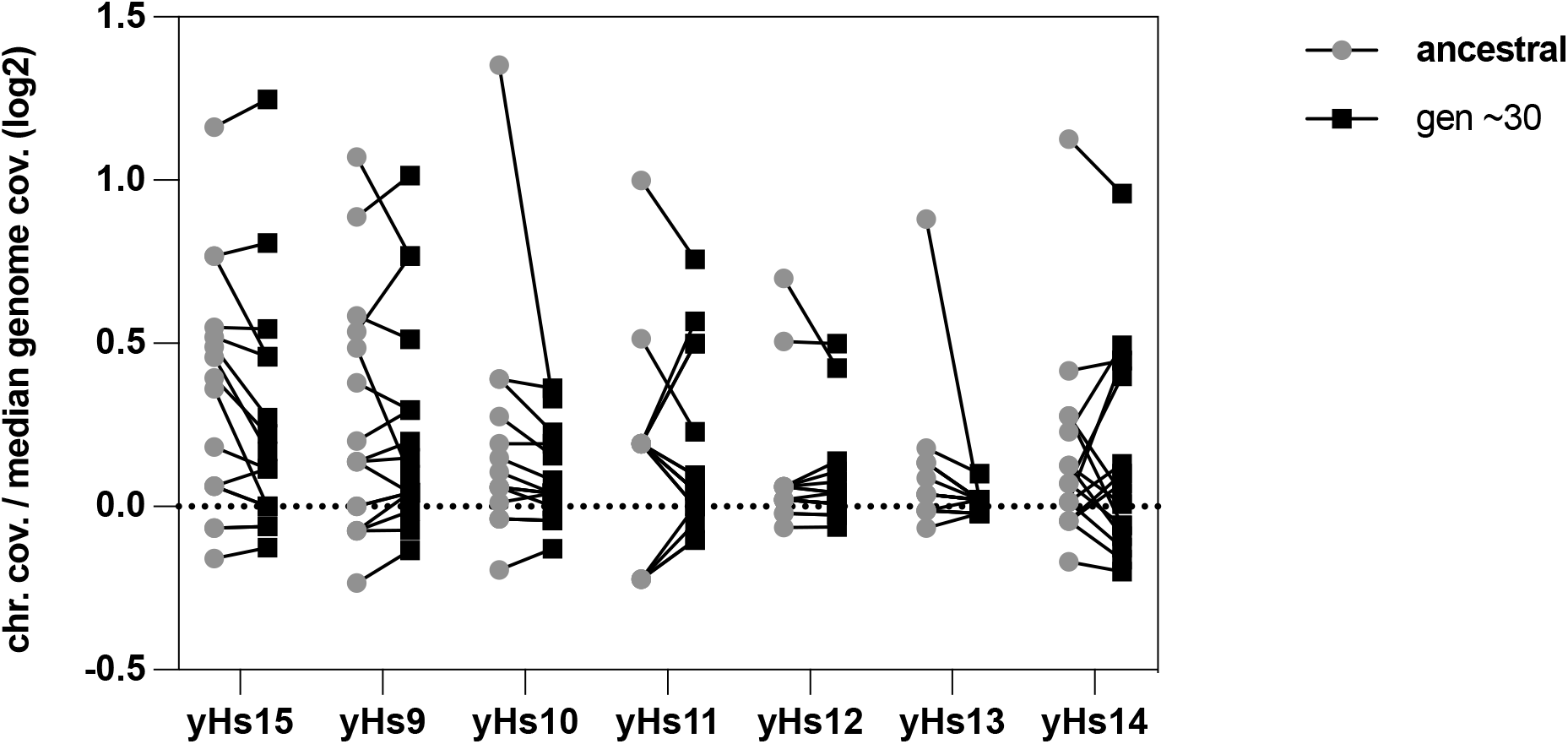
Chromosome aneuploidy evolution in yHs lineages. For each histone–humanized lineage (ancestral, gray circles; and evolved, black squares) we plot the median log2 coverage ratio (chromosome divided by median of genome) for each chromosome.

**Figure S5.**
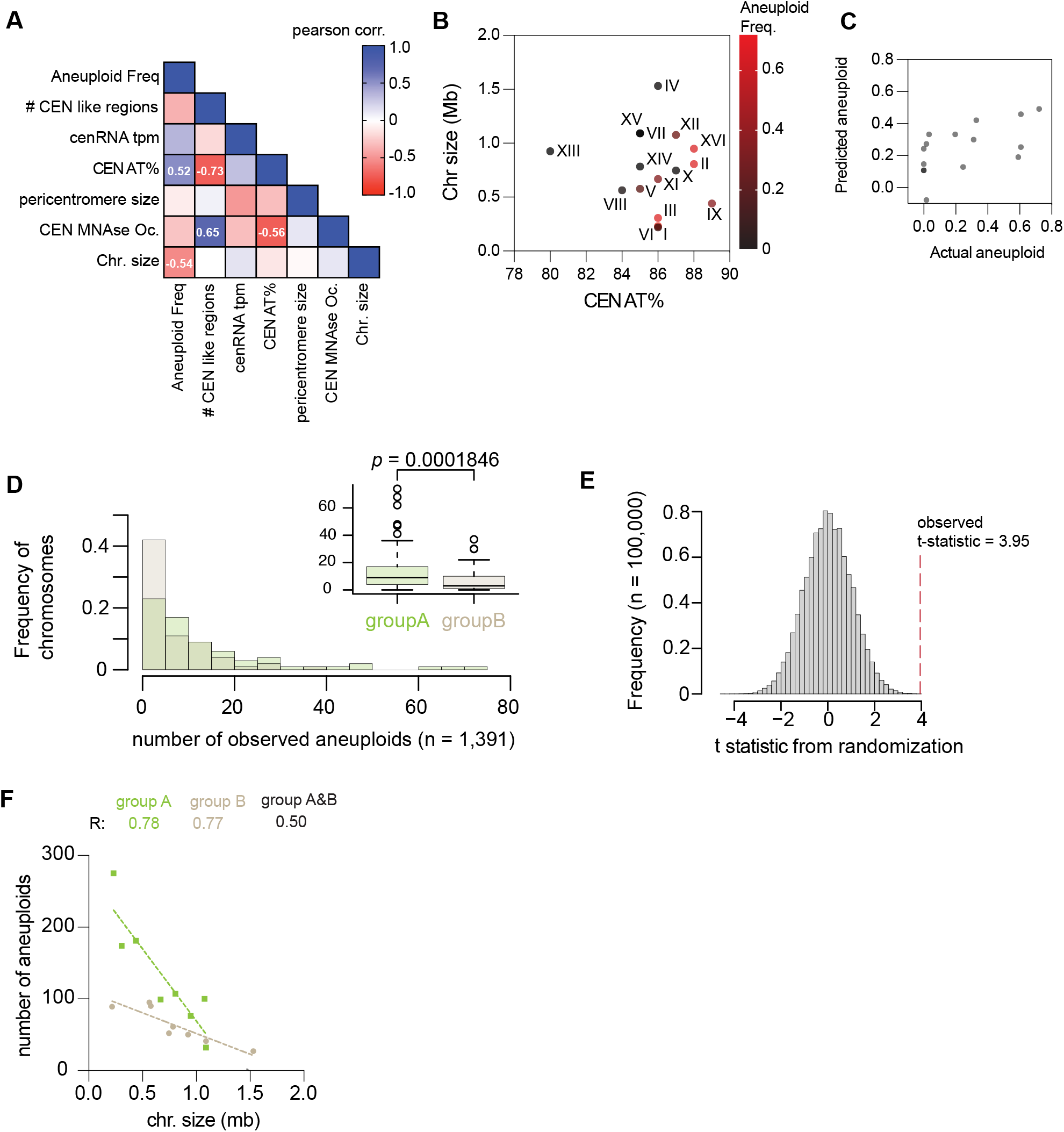
Chromosome size alone does not explain the frequency of aneuploidy in yeasts. (**a**) Correlation matrix of features of centromeres and aneuploidy frequency. Only correlations with significance (p < 0.05) are shown. The number of CEN like regions are taken from Lefrançois et al. (2013) and the pericentromeric sizes are taken from Paldi et al. (2020). (**b**) Plot of centromere AT% as function of chromosome size with the aneuploidy frequency colored in for each point. (**c**) A multiple linear regression model of aneuploidy frequency was constructed using centromere AT% and chromosome size as regression coefficients. Adjusted R^2^ = 0.4188. (**d**) Histogram of the number of aneuploidies per chromosome from this study and seven additional studies (see text). Inset, shows boxplot of the same data with paired t-test of the mean difference in aneuploid frequency between group A and B. (**e**) Histogram of the t-statistics from 100,000 randomized allocations of aneuploid counts between centromeric paralog pairs. The observed t-statistic is shown with a red dashed line. (**f**) Number of observed aneuploidies for each chromosome as a function of chromosome size. R squared values of linear regressions are shown under three models (only group A chromosomes, green; only group B chromosomes, beige; all chromosomes, black). The data is best explained by two models, extra sum-of-squares F test *p* < 0.001.

**Figure S6.**
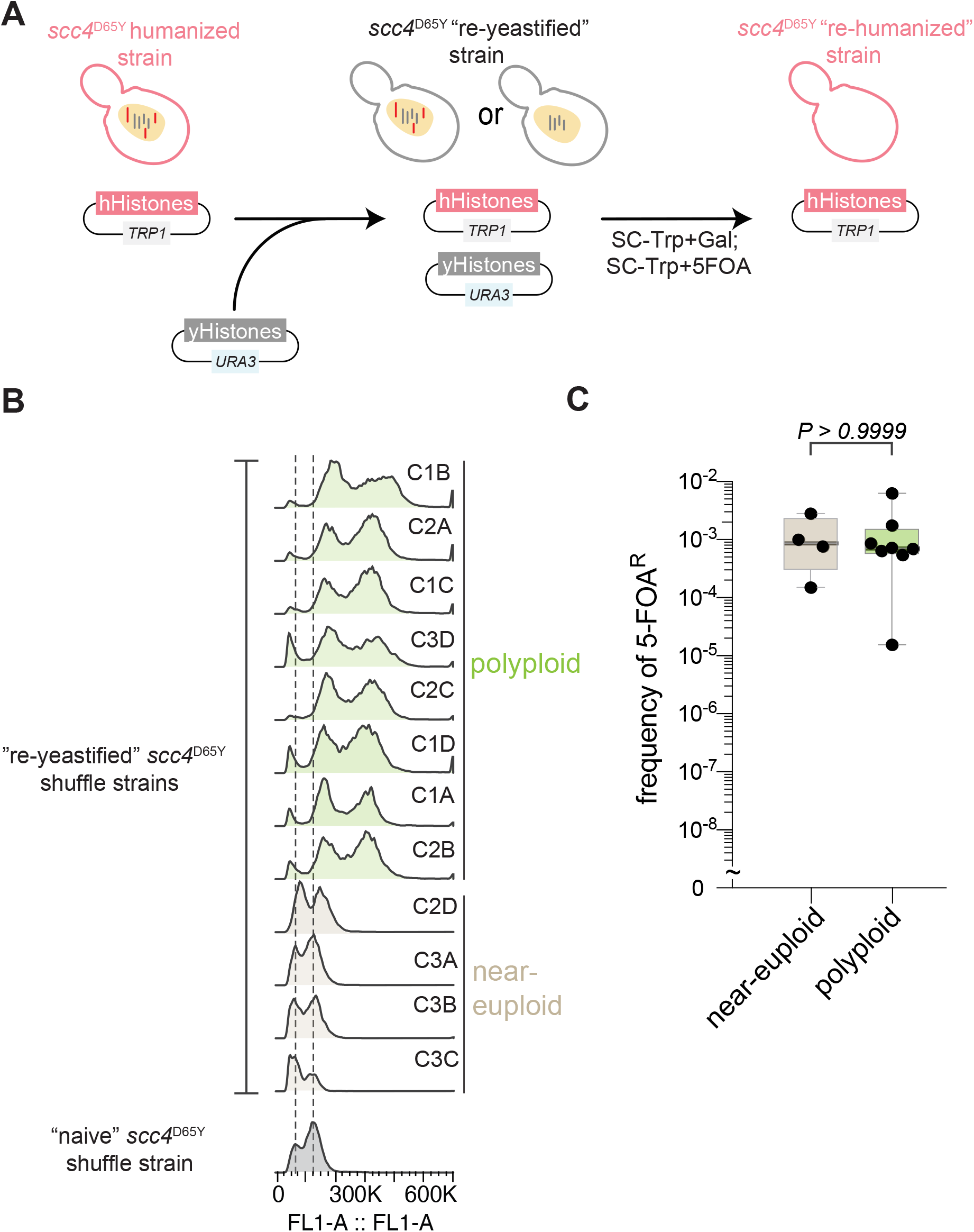
Aneuploidy is non-adaptive for histone-humanization. (**a**) Schematic of the re-yeastification process. First an already histone–humanized strain is transformed with the plasmid encoding all four yeast core histones, then is immediately re-humanized by counterselection of the same. (**b**) Ploidy analysis using flow cytometry of the re-yeastified scc4^D65Y^ strains and the parental strain that has never had human histones. (**c**) Frequency of 5–FOA^R^ resistant colonies following histone–humanization of the re-yeastified strains. Kruskal-Wallis test with Dunn’s multiple corrections; test against the parent scc4^D65Y^ strain show that both euploid and polyploidy strains humanize at significantly higher rates, *p* = 0.009 and *p* = 0.0034, respectively.

**Figure S7.**
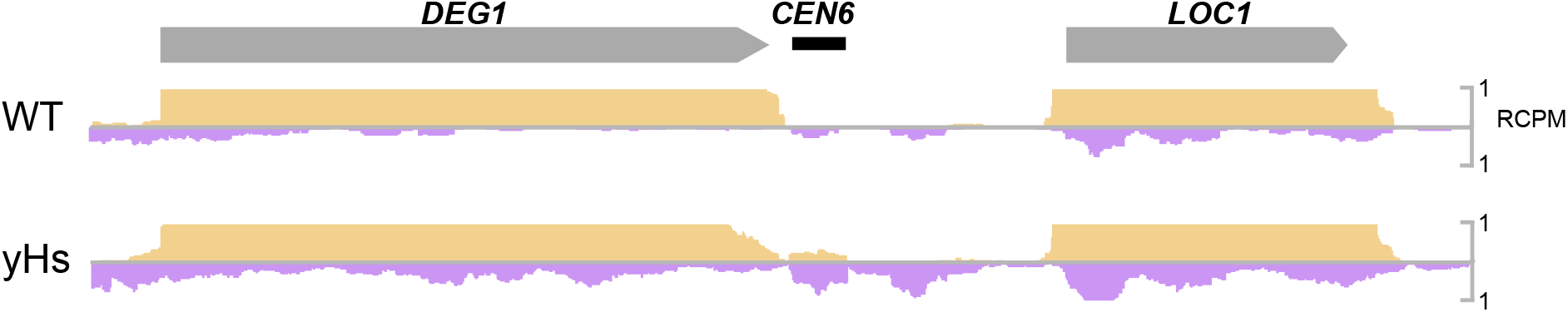
Transcription at chromosomal CEN6 in wild type and histone–humanized yeasts. (**a**) The region around centromere VI is shown, with read coverage of the forward (yellow) and reverse (purple) strands shown.

**Figure S8.**
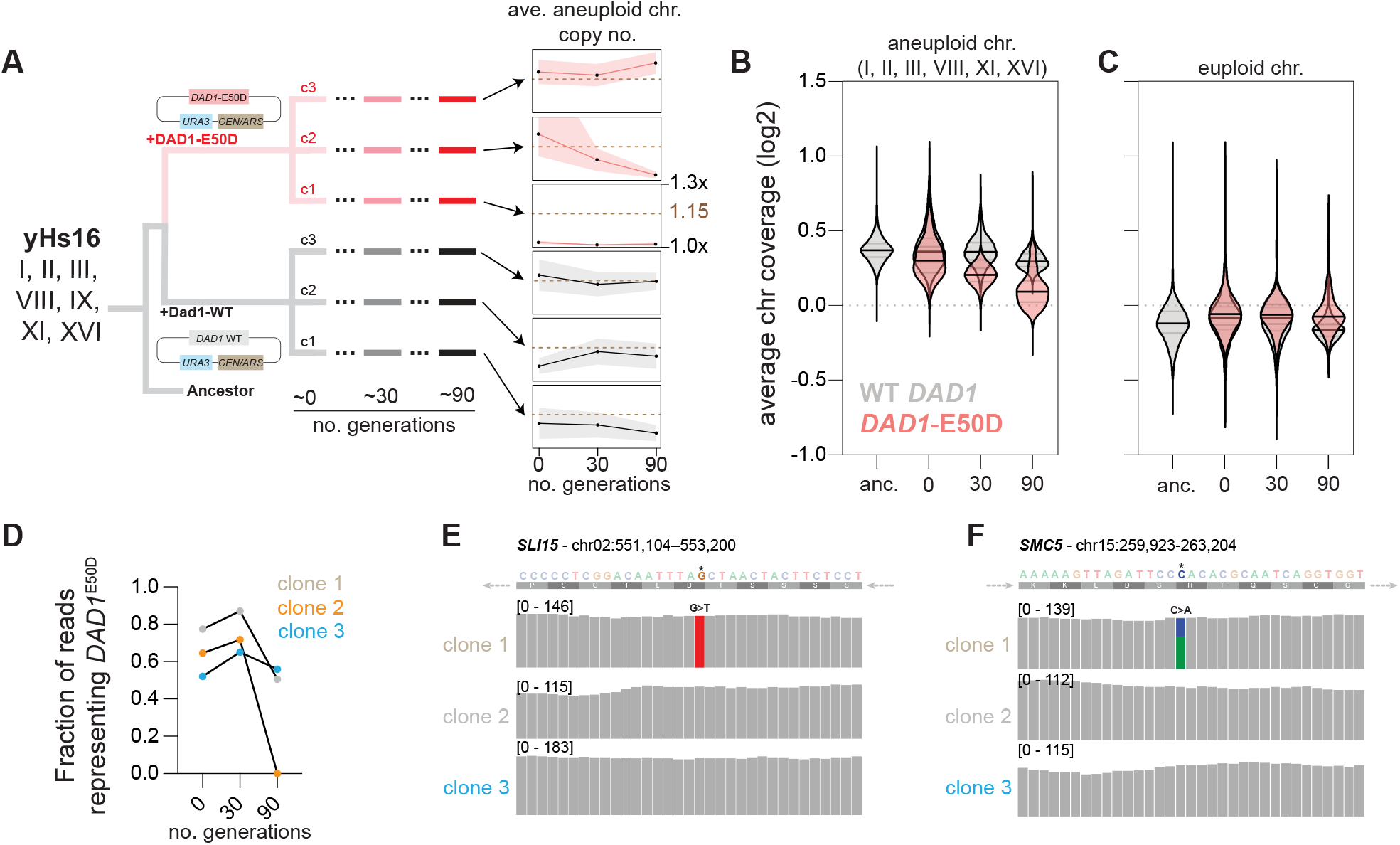
*DAD1*^E50D^ drives euploidization in a heterozygous state. (**a**) Overview of experimental design and ploidy evolution for each clone. Left, histone–humanized clone yH16 was transformed with a CEN/ARS plasmids encoding either the wild type *DAD1* gene or mutant *DAD1*^E50D^ gene. Three clones were selected and passaged for 90 generations in SC–Ura, to maintain selection of the CEN/ARS plasmid. Genomic DNA was isolated and sequenced to infer chromosome copy numbers. Right, plots of the average chromosome copy number for the ancestral aneuploid chromosomes (*I, II, III, VIII, IX, XI*, and *XVI)* at each time point (shaded area represents the standard deviation between the three biological replicates). (**b**) Log2 ratio (chromosome divided by the genome median) chromosomal coverage for the parental aneuploid chromosomes. (**c**) Same as in panel b, however shown for the parental euploid chromosomes. (**d**) Ratio of reads corresponding to the mutant *DAD1*^E50D^ allele for clones transformed with the pRS416-DAD1^E50D^ plasmid across each timepoint. (**e**) Genome browser track for the SLI15 gene, the DNA shown is the Crick strand and represents the template strand of SLI15 (note the arrow of the direction indicates the orientation of SLI15) (**f**) Genome browser track for the SMC5 gene.

**Figure S9.**
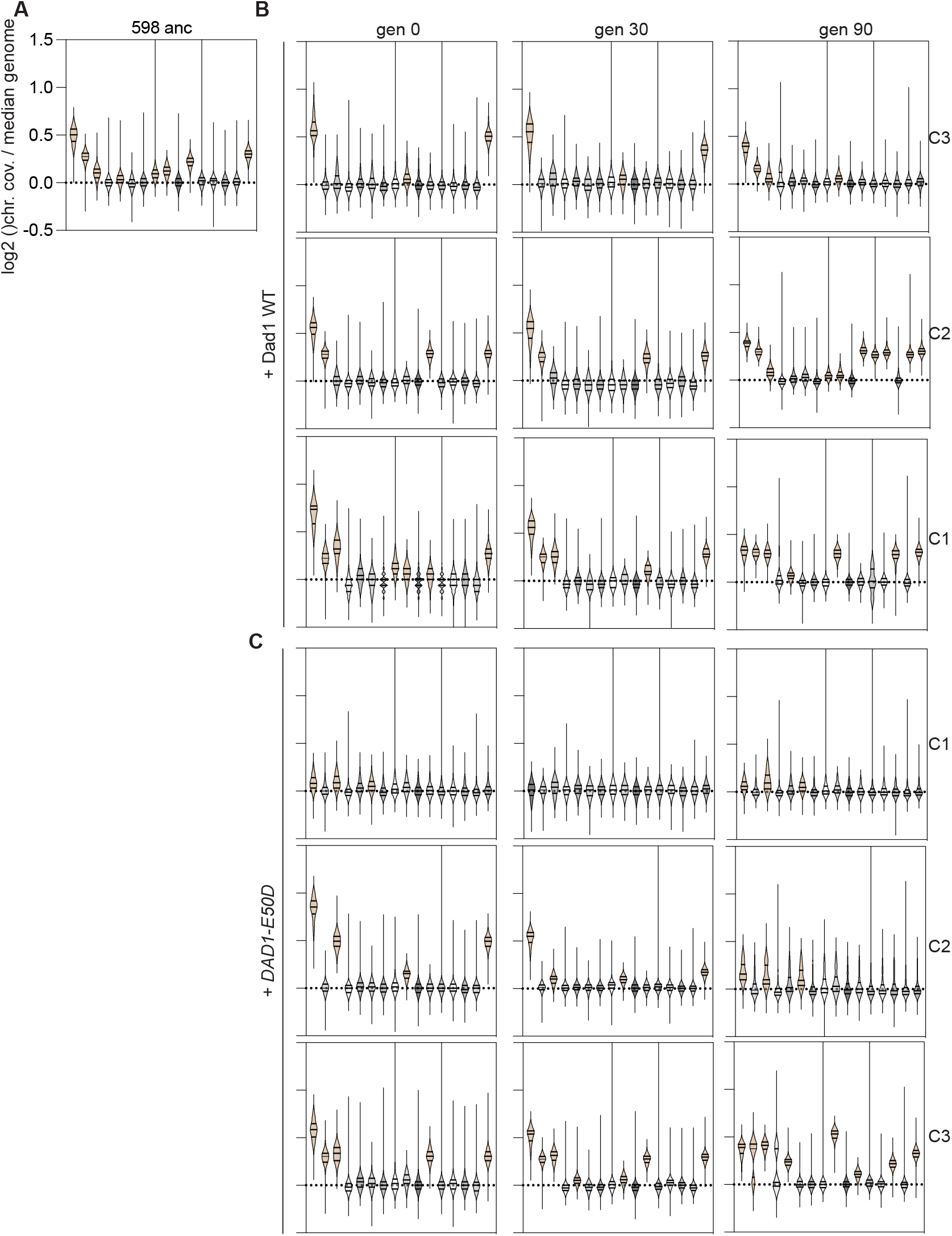
Chromosome copy number of each clone from aneuploidy reduction experiment. (**a**) Chromosome coverage for the ancestral strain prior to transformation with the CEN/ARS plasmid. Violin plots shaded brown are indicated to be aneuploid. (**b**) Chromosome coverage for the clones transformed with wild type *DAD1*. (**c**) Chromosome coverage for the clones transformed with mutant *DAD1*^E50D^.

**Figure S10.**
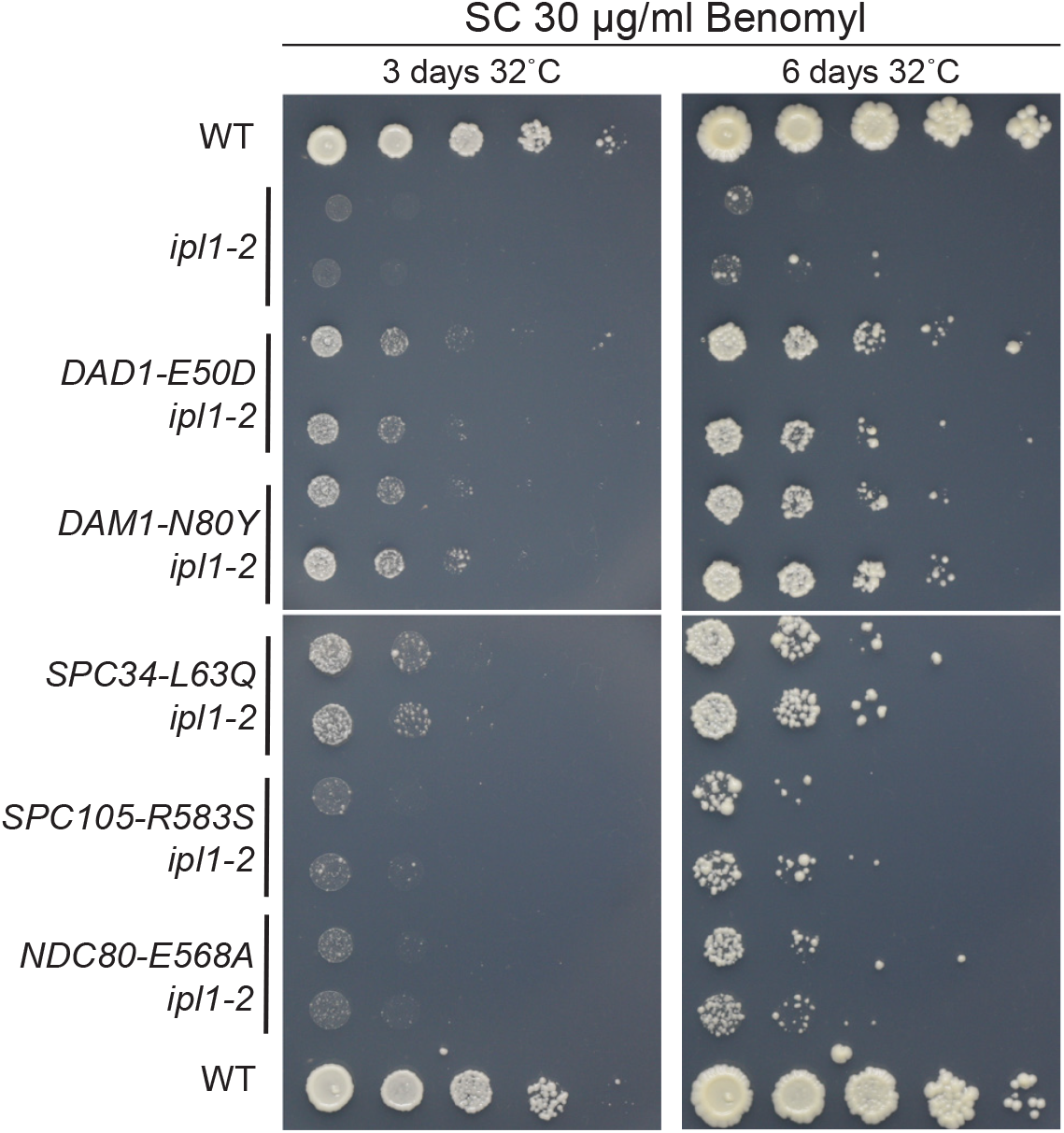
DASH/Dam1c mutants rescue Aurora B kinase ipl1-2 mutant. Growth assays of suppressor mutants with the ipl1-2 temperature sensitive mutant on SC medium with 30 μg/mL of benomyl.

**Figure S11.**
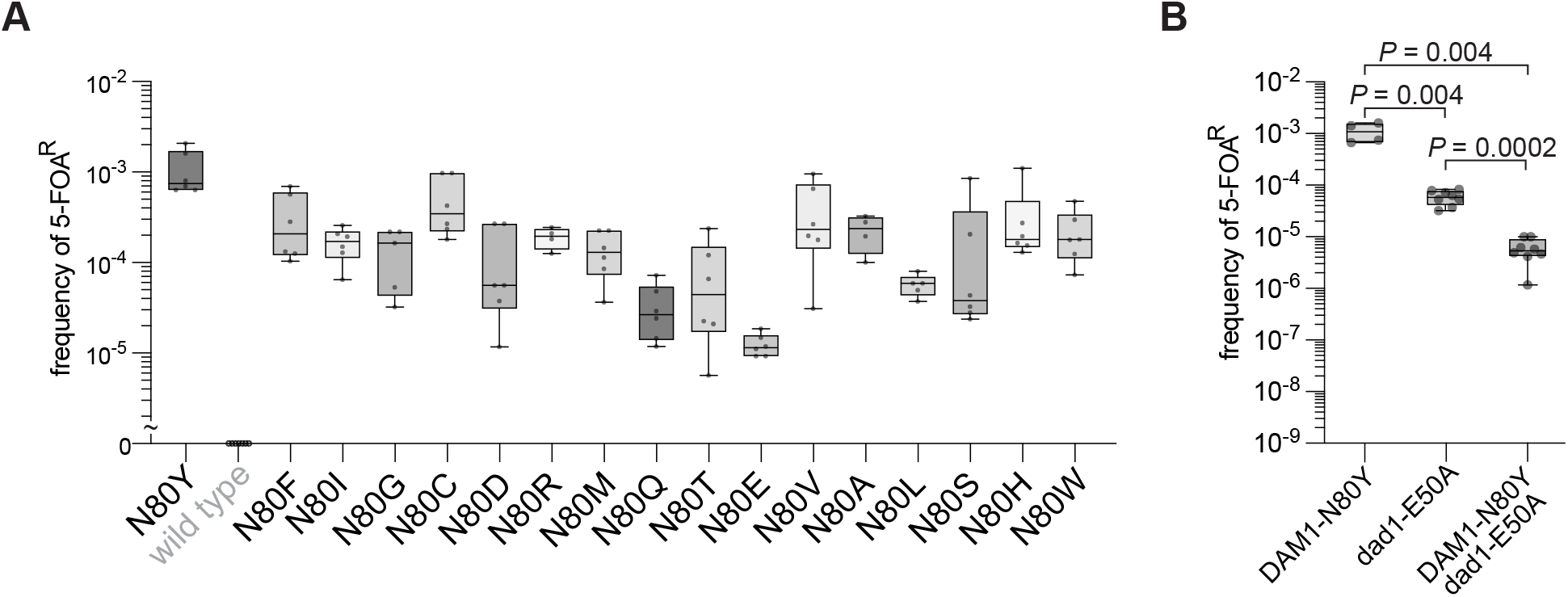
Humanization rates of dam1 residue 80 mutants and *DAM1*^N80Y^ *dad1*^E50A^ mutant. (**a**) Humanization rates (5–FOA^R^ frequency) for the missense mutants of dam1 residue 80. (**b**) Humanization rates (5–FOA^R^ frequency) of the single and double mutants *dad1*^E50A^ and *DAM1*^N80Y^.

**Figure S12.**
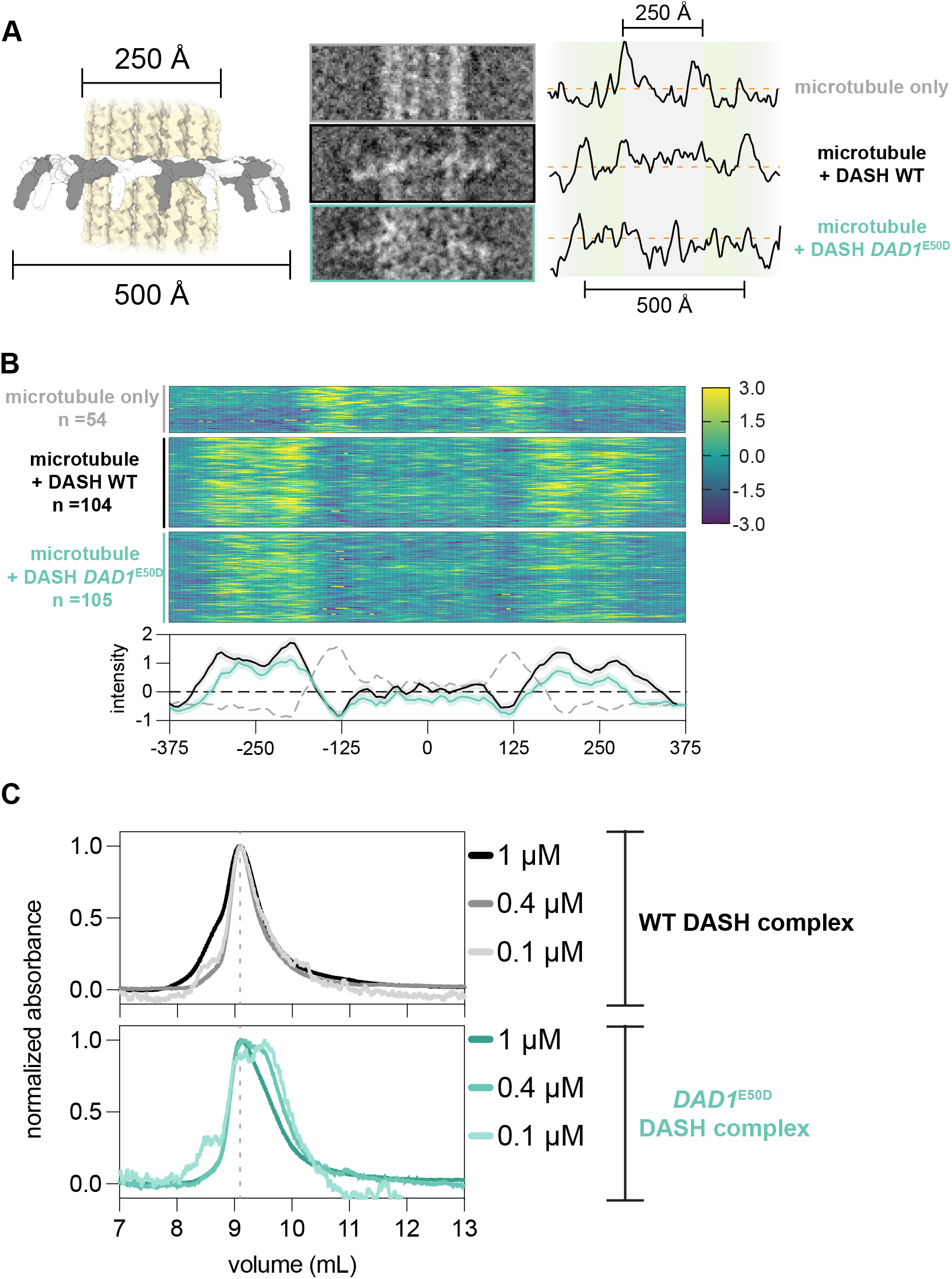
*DAD1*^E50D^ disrupts DASH/Dam1c oligomerization and reduces its dimerization. (**a**) Analysis of DASH complex rings around microtubules. Example images are shown for the three cases measured (bare microtubules, WT DASH, and *DAD1*^E50D^ DASH). A schematic is shown to the left to illustrate the structure of the DASH complex, 6cfz modeled onto a microtubule lattice, 5syf). Note, the bare microtubule lattice was measured from the same micrographs with DASH complex present, with sections showing no complex being chosen to measure. To the right we show the z-score normalized pixel intensities for the example EM micrographs. Note the increased intensity outside the range of the diameter of the microtubule lattice for samples with DASH complex. (**b**) Heatmaps show the z-score normalized pixel intensities along a straight edge bisecting the width of the microtubule or microtubule + DASH complex. The number of particles analyzed is shown. Below are the average profiles of pixel intensity with 95% CI. Dashed gray line is the average profile of pixel intensity of bare microtubules. (**c**) Size exclusion chromatography of WT (upper graph) and mutant *DAD1^E50^D* (lower graph) DASH complexes. Three dilutions of DASH complex are shown. SEC analysis at lower concentrations of the mutant complex, but not for the WT complex, reveal the emergence of a second peak due to a species with a lower apparent molecular weight.

**Figure S13.**
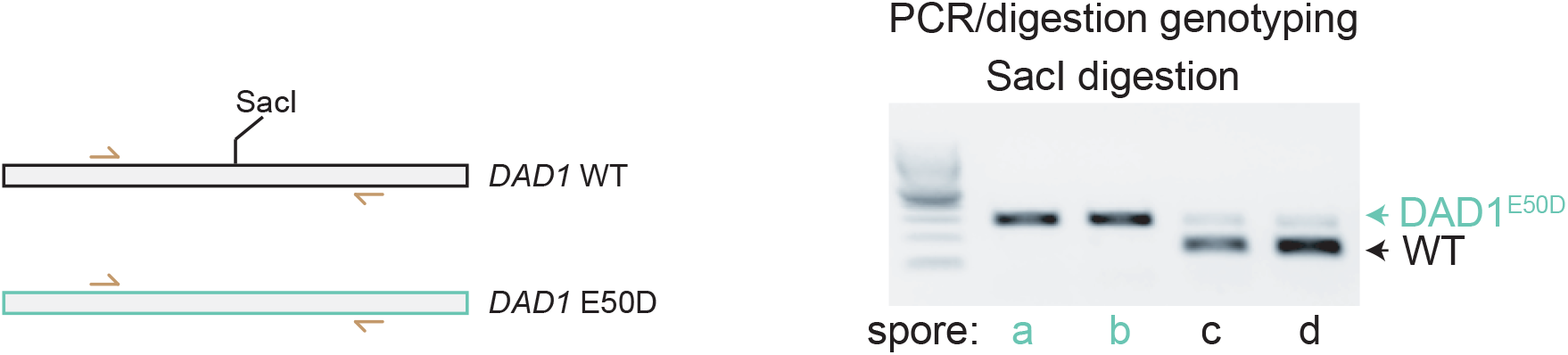
PCR genotyping of *DAD1* in spores from heterozygous cross. (**a**) Schematic and example of the PCR digestion genotype assay used. Briefly, the *DAD1* locus is PCR amplified and then digested with the SacI enzyme, which only cuts the wild type allele. Digestions are then visualized on a 1% agarose gel.

## Bibliography

Abad, M. A., Ruppert, J. G., Buzuk, L., Wear, M., Zou, J., Webb, K. M., Kelly, D. A., Voigt, P., Rappsilber, J., Earnshaw, W. C., and Jeyaprakash, A. A. Borealin–nucleosome interaction secures chromosome association of the chromosomal passenger complex. Journal of Cell Biology, 218(12):3912–3925, Dec. 2019. doi: 10.1083/jcb.201905040.

Akiyoshi, B., Sarangapani, K. K., Powers, A. F., Nelson, C. R., Reichow, S. L., Arellano-Santoyo, H., Gonen, T., Ranish, J. A., Asbury, C. L., and Biggins, S. Tension directly stabilizes reconstituted kinetochore-microtubule attachments. Nature, 468(7323):576–579, Nov. 2010. doi: 10.1038/nature09594.

Antonarakis, S. E. Down syndrome and the complexity of genome dosage imbalance. Nature Reviews Genetics, 18(3):147–163, Mar. 2017. doi: 10.1038/nrg.2016.154.

Asbury, C. L., Gestaut, D. R., Powers, A. F., Franck, A. D., and Davis, T. N. The Dam1 kinetochore complex harnesses microtubule dynamics to produce force and movement. Proceedings of the National Academy of Sciences, 103(26):9873–9878, June 2006. doi: 10.1073/pnas.0602249103.

Baca, S., Prandi, D., Lawrence, M., Mosquera, J., Romanel, A., Drier, Y., Park, K., Kitabayashi, N., MacDonald, T., Ghandi, M., Van Allen, E., Kryukov, G., Sboner, A., Theurillat, J.-P., Soong, T., Nickerson, E., Auclair, D., Tewari, A., Beltran, H., Onofrio, R., Boysen, G., Guiducci, C., Barbieri, C., Cibulskis, K., Sivachenko, A., Carter, S., Saksena, G., Voet, D., Ramos, A., Winckler, W., Cipicchio, M., Ardlie, K., Kantoff, P., Berger, M., Gabriel, S., Golub, T., Meyerson, M., Lander, E., Elemento, O., Getz, G., Demichelis, F., Rubin, M., and Garraway, L. Punctuated Evolution of Prostate Cancer Genomes. Cell, 153(3):666–677, Apr. 2013. doi: 10.1016/j.cell.2013.03.021.

Biggins, S., Severin, F. F., Bhalla, N., Sassoon, I., Hyman, A. A., and Murray, A. W. The conserved protein kinase Ipl1 regulates microtubule binding to kinetochores in budding yeast. Genes & Development, 13(5):532–544, Mar. 1999. doi: 10.1101/gad.13.5.532.

Biggins, S. The Composition, Functions, and Regulation of the Budding Yeast Kinetochore. Genetics, 194(4):817–846, Aug. 2013. doi: 10.1534/genetics.112.145276.

Boonekamp, F. J., Knibbe, E., Vieira-Lara, M. A., Wijsman, M., Luttik, M. A., van Eunen, K., Ridder, M. d., Bron, R., Almonacid Suarez, A. M., van Rijn, P., Wolters, J. C., Pabst, M., Daran, J.-M., Bakker, B. M., and Daran-Lapujade, P. Full humanization of the glycolytic pathway in Saccharomyces cerevisiae. Cell Reports, 39(13):111010, June 2022. doi: 10.1016/j.celrep.2022.111010.

Carmena, M., Wheelock, M., Funabiki, H., and Earnshaw, W. C. The chromosomal passenger complex (CPC): from easy rider to the godfather of mitosis. Nature Reviews Molecular Cell Biology, 13(12):789–803, Dec. 2012. doi: 10.1038/nrm3474.

Chan, C. S. and Botstein, D. Isolation and characterization of chromosome-gain and increase-in-ploidy mutants in yeast. Genetics, 135(3):677–691, Nov. 1993. doi: 10.1093/genetics/135.3.677.

Chen, K., Xi, Y., Pan, X., Li, Z., Kaestner, K., Tyler, J., Dent, S., He, X., and Li, W. DAN-POS: Dynamic analysis of nucleosome position and occupancy by sequencing. Genome Research, 23(2):341–351, Feb. 2013. doi: 10.1101/gr.142067.112.

Cimini, D., Wan, X., Hirel, C. B., and Salmon, E. Aurora Kinase Promotes Turnover of Kinetochore Microtubules to Reduce Chromosome Segregation Errors. Current Biology, 16 (17):1711–1718, Sept. 2006. doi: 10.1016/j.cub.2006.07.022.

Clarke, M. N., Marsoner, T., Alonso Y Adell, M., Ravichandran, M. C., and Campbell, C. S. Multiple routes of adaptation to high levels of CIN and aneuploidy in budding yeast. preprint, bioRxiv, Apr. 2022. URL http://biorxiv.org/lookup/doi/10.1101/2022.04.21.489003.

Cottarel, G., Shero, J. H., Hieter, P., and Hegemann, J. H. A 125-base-pair CEN6 DNA fragment is sufficient for complete meiotic and mitotic centromere functions in Saccharomyces cerevisiae. Molecular and Cellular Biology, 9(8):3342–3349, Aug. 1989. doi: 10.1128/mcb.9.8.3342-3349.1989.

Counter, C. M., Avilion, A. A., LeFeuvre, C. E., Stewart, N. G., Greider, C. W., Harley, C. B., and Bacchetti, S. Telomere shortening associated with chromosome instability is arrested in immortal cells which express telomerase activity. The EMBO journal, 11(5):1921–1929, May 1992. doi: 10.1002/j.1460-2075.1992.tb05245.x.

Duan, S.-F., Han, P.-J., Wang, Q.-M., Liu, W.-Q., Shi, J.-Y., Li, K., Zhang, X.-L., and Bai, F.-Y. The origin and adaptive evolution of domesticated populations of yeast from Far East Asia. Nature Communications, 9(1):2690, Dec. 2018. doi: 10.1038/s41467-018-05106-7.

Flores, R. L., Peterson, Z. E., Zelter, A., Riffle, M., Asbury, C. L., and Davis, T. N. Three interacting regions of the Ndc80 and Dam1 complexes support microtubule tip-coupling under load. Journal of Cell Biology, 221 (5):e202107016, May 2022. doi: 10.1083/jcb.202107016.

Franck, A. D., Powers, A. F., Gestaut, D. R., Gonen, T., Davis, T. N., and Asbury, C. L. Tension applied through the Dam1 complex promotes microtubule elongation providing a direct mechanism for length control in mitosis. Nature Cell Biology, 9(7):832–837, July 2007. doi: 10.1038/ncb1609.

Furuyama, S. and Biggins, S. Centromere identity is specified by a single centromeric nucleosome in budding yeast. Proceedings of the National Academy of Sciences of the United States of America, 104(37):14706–14711, Sept. 2007. doi: 10.1073/pnas.0706985104.

Gallone, B., Steensels, J., Prahl, T., Soriaga, L., Saels, V., Herrera-Malaver, B., Merlevede, A., Roncoroni, M., Voordeckers, K., Miraglia, L., Teiling, C., Steffy, B., Taylor, M., Schwartz, A., Richardson, T., White, C., Baele, G., Maere, S., and Verstrepen, K. J. Domestication and Divergence of Saccharomyces cerevisiae Beer Yeasts. Cell, 166(6):1397–1410.e16, Sept. 2016. doi: 10.1016/j.cell.2016.08.020.

Gordon, J. L., Byrne, K. P., and Wolfe, K. H. Mechanisms of Chromosome Number Evolution in Yeast. PLoS Genetics, 7(7):e1002190, July 2011. doi: 10.1371/journal.pgen.1002190.

Gould, S. j. and Eldredge, N. Punctuated equilibrium comes of age. Nature, 366(6452): 223–227, Nov. 1993. doi: 10.1038/366223a0.

Haase, M. A., Truong, D. M., and Boeke, J. D. Superloser: a plasmid shuffling vector for *Saccharomyces cerevisiae* with exceedingly low background. G3 Genes|Genomes|Genetics, 9, Aug. 2019. doi: 10.1534/g3.119.400325.

Heasley, L. R., Sampaio, N. M. V., and Argueso, J. L. Systemic and rapid restructuring of the genome: a new perspective on punctuated equilibrium. Current Genetics, 67(1):57–63, Feb. 2021. doi: 10.1007/s00294-020-01119-2.

Hedouin, S., Logsdon, G. A., Underwood, J. G., and Biggins, S. A transcriptional roadblock protects yeast centromeres. Nucleic Acids Research, page gkac117, Mar. 2022. doi: 10.1093/nar/gkac117.

Hose, J., Escalante, L. E., Clowers, K. J., Dutcher, H. A., Robinson, D., Bouriakov, V., Coon, J. J., Shishkova, E., and Gasch, A. P. The genetic basis of aneuploidy tolerance in wild yeast. eLife, 9:e52063, Jan. 2020. doi: 10.7554/eLife.52063.

Jenni, S. and Harrison, S. C. Structure of the DASH/Dam1 complex shows its role at the yeast kinetochore-microtubule interface. Science, 360(6388):552–558, May 2018. doi: 10.1126/science.aar6436.

Jenni, S., Dimitrova, Y. N., Valverde, R., Hinshaw, S. M., and Harrison, S. C. Molecular Structures of Yeast Kinetochore Subcomplexes and Their Roles in Chromosome Segregation. Cold Spring Harbor Symposia on Quantitative Biology, 82:83–89, 2017. doi: 10.1101/sqb.2017.82.033738.

Kachroo, A. H., Laurent, J. M., Yellman, C. M., Meyer, A. G., Wilke, C. O., and Marcotte, E. M. Systematic humanization of yeast genes reveals conserved functions and genetic modularity. Science, 348(6237):921–925, May 2015. doi: 10.1126/science.aaa0769.

Kao, K. C., Schwartz, K., and Sherlock, G. A Genome-Wide Analysis Reveals No Nuclear Dobzhansky-Muller Pairs of Determinants of Speciation between S. cerevisiae and S. paradoxus, but Suggests More Complex Incompatibilities. PLoS Genetics, 6(7): e1001038, July 2010. doi: 10.1371/journal.pgen.1001038.

Kawashima, S. A., Yamagishi, Y., Honda, T., Ishiguro, K.-i., and Watanabe, Y. Phosphorylation of H2A by Bub1 Prevents Chromosomal Instability Through Localizing Shugoshin. Science, 327(5962):172–177, Jan. 2010. doi: 10.1126/science.1180189.

Kim, J. o., Zelter, A., Umbreit, N. T., Bollozos, A., Riffle, M., Johnson, R., MacCoss, M. J., Asbury, C. L., and Davis, T. N. The Ndc80 complex bridges two Dam1 complex rings. eLife, 6:e21069, Feb. 2017. doi: 10.7554/eLife.21069.

Laurent, J. M., Garge, R. K., Teufel, A. I., Wilke, C. O., Kachroo, A. H., and Marcotte, E. M. Humanization of yeast genes with multiple human orthologs reveals functional divergence between paralogs. PLOS Biology, 18(5):e3000627, May 2020. doi: 10.1371/journal.pbio.3000627.

Ling, Y. H. and Yuen, K. W. Y. Point centromere activity requires an optimal level of centromeric noncoding RNA. Proceedings of the National Academy of Sciences, 116(13): 6270–6279, Mar. 2019. doi: 10.1073/pnas.1821384116.

Marcet-Houben, M. and Gabaldón, T. Beyond the Whole-Genome Duplication: Phylogenetic Evidence for an Ancient Interspecies Hybridization in the Baker’s Yeast Lineage. PLOS Biology, 13(8):e1002220, Aug. 2015. doi: 10.1371/journal.pbio.1002220.

McCulley, J. L. and Petes, T. D. Chromosome rearrangements and aneuploidy in yeast strains lacking both Tel1p and Mec1p reflect deficiencies in two different mechanisms. Proceedings of the National Academy of Sciences, 107(25):11465–11470, June 2010. doi: 10.1073/pnas.1006281107.

Miranda, J. L., Wulf, P. D., Sorger, P. K., and Harrison, S. C. The yeast DASH complex forms closed rings on microtubules. Nature Structural & Molecular Biology, 12(2):138–143, Feb. 2005. doi: 10.1038/nsmb896.

Musacchio, A. and Salmon, E. D. The spindle-assembly checkpoint in space and time. Nature Reviews Molecular Cell Biology, 8(5):379–393, May 2007. doi: 10.1038/nrm2163.

Nicklas, R. B. How Cells Get the Right Chromosomes. Science, 275(5300):632–637, Jan. 1997. doi: 10.1126/science.275.5300.632.

Oromendia, A. B. and Amon, A. Aneuploidy: implications for protein homeostasis and disease. Disease Models & Mechanisms, 7(1):15–20, Jan. 2014. doi: 10.1242/dmm.013391.

Peter, J., De Chiara, M., Friedrich, A., Yue, J.-X., Pflieger, D., Bergström, A., Sigwalt, A., Barre, B., Freel, K., Llored, A., Cruaud, C., Labadie, K., Aury, J.-M., Istace, B., Lebrigand, K., Barbry, P., Engelen, S., Lemainque, A., Wincker, P., Liti, G., and Schacherer, J. Genome evolution across 1,011 Saccharomyces cerevisiae isolates. Nature, 556(7701): 339–344, Apr. 2018. doi: 10.1038/s41586-018-0030-5.

Pinsky, B. A., Kung, C., Shokat, K. M., and Biggins, S. The Ipl1-Aurora protein kinase activates the spindle checkpoint by creating unattached kinetochores. Nature Cell Biology, 8 (1):78–83, Jan. 2006. doi: 10.1038/ncb1341.

Powers, A. F., Franck, A. D., Gestaut, D. R., Cooper, J., Gracyzk, B., Wei, R. R., Wordeman, L., Davis, T. N., and Asbury, C. L. The Ndc80 Kinetochore Complex Forms Load-Bearing Attachments to Dynamic Microtubule Tips via Biased Diffusion. Cell, 136(5):865–875, Mar. 2009. doi: 10.1016/j.cell.2008.12.045.

Sarangapani, K. K., Duro, E., Deng, Y., Alves, F. d. L., Ye, Q., Opoku, K. N., Ceto, S., Rappsilber, J., Corbett, K. D., Biggins, S., Marston, A. L., and Asbury, C. L. Sister kinetochores are mechanically fused during meiosis I in yeast. Science, 346(6206):248–251, Oct. 2014. doi: 10.1126/science.1256729.

Schindelin, J., Arganda-Carreras, I., Frise, E., Kaynig, V., Longair, M., Pietzsch, T., Preibisch, S., Rueden, C., Saalfeld, S., Schmid, B., Tinevez, J.-Y., White, D. J., Hartenstein, V., Eliceiri, K., Tomancak, P., and Cardona, A. Fiji: an open-source platform for biological-image analysis. Nature Methods, 9(7):676–682, July 2012. doi: 10.1038/nmeth.2019.

Sharp, N. P., Sandell, L., James, C. G., and Otto, S. P. The genome-wide rate and spectrum of spontaneous mutations differ between haploid and diploid yeast. Proceedings of the National Academy of Sciences, 115(22), May 2018. doi: 10.1073/pnas.1801040115.

Shen, X.-X., Opulente, D. A., Kominek, J., Zhou, X., Steenwyk, J. L., Buh, K. V., Haase, M. A., Wisecaver, J. H., Wang, M., Doering, D. T., Boudouris, J. T., Schneider, R. M., Langdon, Q. K., Ohkuma, M., Endoh, R., Takashima, M., Manabe, R.-i., Čadež, N., Libkind, D., Rosa, C. A., DeVirgilio, J., Hulfachor, A. B., Groenewald, M., Kurtzman, C. P., Hittinger, C. T., and Rokas, A. Tempo and Mode of Genome Evolution in the Budding Yeast Subphylum. Cell, 175(6):1533–1545.e20, Nov. 2018. doi: 10.1016/j.cell.2018.10.023.

Steiner, F. A. and Henikoff, S. Diversity in the organization of centromeric chromatin. Current Opinion in Genetics & Development, 31:28–35, Apr. 2015. doi: 10.1016/j.gde.2015.03.010.

Tanaka, T. U., Rachidi, N., Janke, C., Pereira, G., Galova, M., Schiebel, E., Stark, M. J., and Nasmyth, K. Evidence that the Ipl1-Sli15 (Aurora Kinase-INCENP) Complex Promotes Chromosome Bi-orientation by Altering Kinetochore-Spindle Pole Connections. Cell, 108 (3):317–329, Feb. 2002. doi: 10.1016/S0092-8674(02)00633-5.

Tien, J. F., Umbreit, N. T., Gestaut, D. R., Franck, A. D., Cooper, J., Wordeman, L., Gonen, T., Asbury, C. L., and Davis, T. N. Cooperation of the Dam1 and Ndc80 kinetochore complexes enhances microtubule coupling and is regulated by aurora B. Journal of Cell Biology, 189(4):713–723, May 2010. doi: 10.1083/jcb.200910142.

Torres, E. M., Sokolsky, T., Tucker, C. M., Chan, L. Y., Boselli, M., Dunham, M. J., and Amon, A. Effects of Aneuploidy on Cellular Physiology and Cell Division in Haploid Yeast. Science, 317(5840):916–924, Aug. 2007. doi: 10.1126/science.1142210.

Torres, E. M., Dephoure, N., Panneerselvam, A., Tucker, C. M., Whittaker, C. A., Gygi, S. P., Dunham, M. J., and Amon, A. Identification of Aneuploidy-Tolerating Mutations. Cell, 143 (1):71–83, Oct. 2010. doi: 10.1016/j.cell.2010.08.038.

Truong, D. M. and Boeke, J. D. Resetting the Yeast Epigenome with Human Nucleosomes. Cell, 171(7):1508–1519.e13, Dec. 2017. doi: 10.1016/j.cell.2017.10.043.

Umbreit, N. T., Miller, M. P., Tien, J. F., Cattin Ortolá, J., Gui, L., Lee, K. K., Biggins, S., Asbury, C. L., and Davis, T. N. Kinetochores require oligomerization of Dam1 complex to maintain microtubule attachments against tension and promote biorientation. Nature Communications, 5(1):4951, Dec. 2014. doi: 10.1038/ncomms5951.

Waterhouse, A., Bertoni, M., Bienert, S., Studer, G., Tauriello, G., Gumienny, R., Heer, F. T., de Beer, T. A., Rempfer, C., Bordoli, L., Lepore, R., and Schwede, T. SWISS-MODEL: homology modelling of protein structures and complexes. Nucleic Acids Research, 46 (W1):W296–W303, July 2018. doi: 10.1093/nar/gky427.

Winey, M., Mamay, C. L., O’Toole, E. T., Mastronarde, D. N., Giddings, T. H., McDonald, K. L., and McIntosh, J. R. Three-dimensional ultrastructural analysis of the Saccharomyces cerevisiae mitotic spindle. Journal of Cell Biology, 129(6):1601–1615, June 1995. doi: 10.1083/jcb.129.6.1601.

Wolfe, K. H. and Shields, D. C. Molecular evidence for an ancient duplication of the entire yeast genome. Nature, 387(6634):708–713, June 1997. doi: 10.1038/42711.

Yamagishi, Y., Honda, T., Tanno, Y., and Watanabe, Y. Two Histone Marks Establish the Inner Centromere and Chromosome Bi-Orientation. Science, 330(6001):239–243, Oct. 2010. doi: 10.1126/science.1194498.

Zaytsev, A. V. and Grishchuk, E. L. Basic mechanism for biorientation of mitotic chromosomes is provided by the kinetochore geometry and indiscriminate turnover of kinetochore microtubules. Molecular Biology of the Cell, 26(22):3985–3998, Nov. 2015. doi: 10.1091/mbc.E15-06-0384.

Zhu, Y. O., Sherlock, G., and Petrov, D. A. Whole Genome Analysis of 132 Clinical *Saccharomyces cerevisiae* Strains Reveals Extensive Ploidy Variation. G3 Genes|Genomes|Genetics, 6(8):2421–2434, Aug. 2016. doi: 10.1534/g3.116.029397.

## Supplemental Bibliography

Lefrançois, P., Auerbach, R. K., Yellman, C. M., Roeder, G. S., and Snyder, M. Centromere-Like Regions in the Budding Yeast Genome. PLoS Genetics, 9(1):e1003209, Jan. 2013. doi: 10.1371/journal.pgen.1003209.

Paldi, F., Alver, B., Robertson, D., Schalbetter, S. A., Kerr, A., Kelly, D. A., Baxter, J., Neale, M. J., and Marston, A. L. Convergent genes shape budding yeast pericentromeres. Nature, 582 (7810):119–123, June 2020. doi: 10.1038/s41586-020-2244-6.

